# Characterization of a GpsB-associated regulator of PBP1a reveals the organization of the cell wall remodeling complex of *Streptococcus pneumoniae*

**DOI:** 10.1101/2024.11.09.622756

**Authors:** Hugo Millat, Cassandra Lenoir, Cassandra Falcou, Caroline Cluzel, André Zapun, David I Roper, Cécile Morlot, Adrien Ducret, Christophe Grangeasse

## Abstract

Class A PBPs (aPBPs) play a key role in the biosynthesis and remodeling of peptidoglycan, the main component of the bacterial cell wall. The human bacterial pathogen *Streptococcus pneumoniae* produces three aPBPs, which are regulated to maintain the bacterium’s ovoid shape. Although their exact functions remain unclear, evidence suggests that PBP1a and PBP2a activities are closely coordinated. In this study, we elucidated the function of an unknown function protein named GarP (<u>G</u>psB-<u>a</u>ssociated regulator of <u>P</u>BP1a), in the regulation of PBP1a activity. We showed that GarP localizes to the division septum and its absence leads to morphological defects. We further identified a GpsB-binding motif in GarP as well as in PBP2a, the PG deacetylase PgdA and the muramidase MpgA. Our analysis of genetic and protein interactions, combined with cell imaging, supports a model of a molecular complex that coordinates PG remodeling during *S. pneumoniae* cell division.

## INTRODUCTION

Peptidoglycan (PG) is a critical component of bacterial cell walls, providing structural integrity and protection from osmotic lysis and cell shape maintenance^1^. It consists of glycan strands made of repeating units of N-acetylglucosamine (GlcNAc) and N-acetylmuramic acid (MurNAc), with short peptide chains attached to the MurNAc that crosslink adjacent glycan strands^2^. This polymer forms a robust, mesh-like polymeric matrix around the cytoplasmic membrane. PG synthesis involves 3 main types of synthases: class A bifunctional penicillin-binding proteins (aPBPs), class B penicillin-binding proteins (bPBPs) and SEDS (Shape, Elongation, Division, and Sporulation) proteins^2^. aPBPs are enzymes with both Lipid II glycosyltransferase (GT) activity for glycan polymerization and transpeptidase (TP) activity for peptide crosslinking of the glycan strand that is produced by the GT site. On the other hand, SEDS proteins provide only the GT activity and bPBPs perform only the TP activity. The glycan strand product of the SEDS protein Lipid II glycosyltransferase activity forms a substrate for the transpeptidase activity of the bPBP. According to the current models and understanding of peptidoglycan biosynthesis, SEDS and bPBPs work together to assemble the PG primary framework, which is further remodeled by the GT/TP activities of aPBPs^3,4^. Thus, overall peptidoglycan biosynthesis requires both SEDS-bPBP and aPBP activities for normal cell growth and division.

The PG primary framework is assembled by two distinct multiprotein machines called the divisome and the elongasome, both of which are essential and function independently to allow cell division and cell elongation, respectively^5^. The divisome localizes at midcell and is organized by the tubulin-like protein FtsZ^6^. It contains the SEDS FtsW, which works as a functionally co-dependent complex with a cognate bPBP, which is FtsI (PBP3) in *Escherichia coli*, PBP2b in *Bacillus subtilis* and PBP2x in *Streptococcus pneumoniae*^7,8^. By contrast, the elongasome, which includes the SEDS-bPBP core enzymatic complex between RodA and PBP2, is localized either along the sidewalls (*E. coli* or *B. subtilis)* or at the poles (*Mycobacterium tuberculosis*) in rod-shaped bacteria and is organized by the actin-like protein MreB^9^ or the tropomyosin-like protein DivIVA, respectively^10^. However, unlike rod- shaped cells, cell elongation of the ovoid-shaped pathogen *S. pneumoniae*, which lacks MreB, takes place at midcell and is also organized by FtsZ^5^ and requires the SEDS-bPBP pair RodA-PBP2b^11^.

The role of the aPBPs are proposed to be the remodeling of the PG layer and/or repair its damaged areas, ensuring that the cell wall remains intact and functional under various stress conditions^4,12^. In Gram-negative bacteria, aPBPs are activated by outer membrane lipoproteins. For instance, PBP1a and PBP1b of *E. coli* are activated by the lipoproteins LpoA and LpoB, respectively^13^. Each lipoprotein interacts with specific regulatory domains (OB/ODD for PBP1a and UB2H for PBP1b), inducing conformational changes and activating their PG polymerization activity^14,15^. In Gram-positive bacteria, the regulation of aPBPs is less understood, but several proteins have been proposed to control their function. Substantial evidence supports the role of scaffolding proteins, such as the DivIVA paralog GpsB, in the spatiotemporal regulation of aPBPs. For instance, the subcellular localization of the aPBPs PBP1, PBPA1 or PBP2 at the septum is determined by their direct interaction with GpsB in *B. subtilis, Listeria monocytogenes* or *Staphylococcus aureus*, respectively^16,17^. This specific interaction involves the consensus GpsB-binding motif [(S/T)RxxR(R/K)] found in the cytoplasmic domain of these PBPs. In the case of *S. pneumoniae*, which produces three different aPBPs (PBP1a, PBP1b, PBP2a)^1^, only PBP2a possesses this GpsB-binding motif^17^ and other regulatory mechanisms also control aPBP localization and/or activity. For example, the membrane proteins CozEa and CopD influence the activity and/or the localization of PBP1a but the mechanism behind this regulation is still unknown^18,19^. To our knowledge, the best described aPBP regulator is MacP, which controls the function of PBP2a^20^. It is proposed that MacP enhances PBP2a activity by altering the interface between the GT domain and the transmembrane helix of PBP2a^21^.

Pneumococcal mutants inactivated for either *pbp1a*, *pbp1b* or *pbp2a* are viable, and only simultaneous inactivation of *pbp1a* and *pbp2a* is lethal, suggesting that they are at least partially functionally redundant^22^. Importantly, both PBP1a and PBP2a interact with GpsB but only PBP2a possesses the consensus GpsB-binding motif [(S/T)RxxR(R/K)]^17^. Moreover, only *pbp1a*, but not *pbp2a*, become essential in the absence of *gpsB*, indicating that PBP2a is controlled by GpsB^23^. Taken together, it is likely that the two major aPBPs in *S. pneumoniae*, PBP1a and PBP2a, fulfill both specific and overlapping functions, and are thus scaffolded and/or regulated by distinct and common proteins to ensure the PG layer is fully matured or repaired. However, how these scaffolding proteins and potential regulators control the localization and/or activity of aPBPs along the cell cycle remains unknown.

In this study, we elucidated the role of a membrane protein of previously unknown function (Spr1457) in regulating PBP1a activity, the major aPBP associated with the pneumococcal elongasome. We named this protein GarP for GpsB-associated regulator of PBP1a. By combining genetics, *in vivo* imaging and biochemical approaches, we demonstrate that GarP promotes the glycosyltransferase activity of PBP1a by a direct interaction. We further show that this activation may result from the stabilization of the catalytic site of PBP1a by a predicted α-helix of GarP. Importantly, we demonstrate that GarP possesses the consensus GpsB-binding motif and interacts directly with GpsB. We further show that this motif is observed in the PG deacetylase PgdA and the muramidase MpgA. Further observations allow us to propose that a complex, composed of GarP/PBP1a/MacP/PBP2a/MpgA/PgdA and coordinated by GpsB, could be specifically responsible for PG remodeling during cell division in *S. pneumoniae*.

## RESULTS

### GarP is a membrane protein that forms a ring at midcell

In an attempt to identify new proteins involved in cell wall assembly in *S. pneumoniae*, we focused on proteins of unknown function that contain a LysM PG-binding domain^24^. By searching the pneumococcal R6 genomic database, we identified 4 proteins with a putative LysM domain. Beside the muramidase MpgA^25^ and two putative PG hydrolases Spr0096 and Spr1875^26^, we identified Spr1457 (henceforth named GarP for GpsB-associated regulator of PBP1a, on the basis of the observations we report here), a protein that contains a single LysM domain and no other predicted functional domain (Fig. 1a). GarP is a 158-amino acid protein with a single predicted transmembrane (TM) domain. Both the N-terminal cytoplasmic region and that between the TM and the LysM domain are predicted to be intrinsically disordered by MobiDB (https://mobidb.org). A high-confidence AlphaFold3 prediction (https://alphafoldserver.com) further suggests that an α-helix (referred to as PαH below) may form after the TM segment (Fig. 1a, b).

**Fig. 1:**
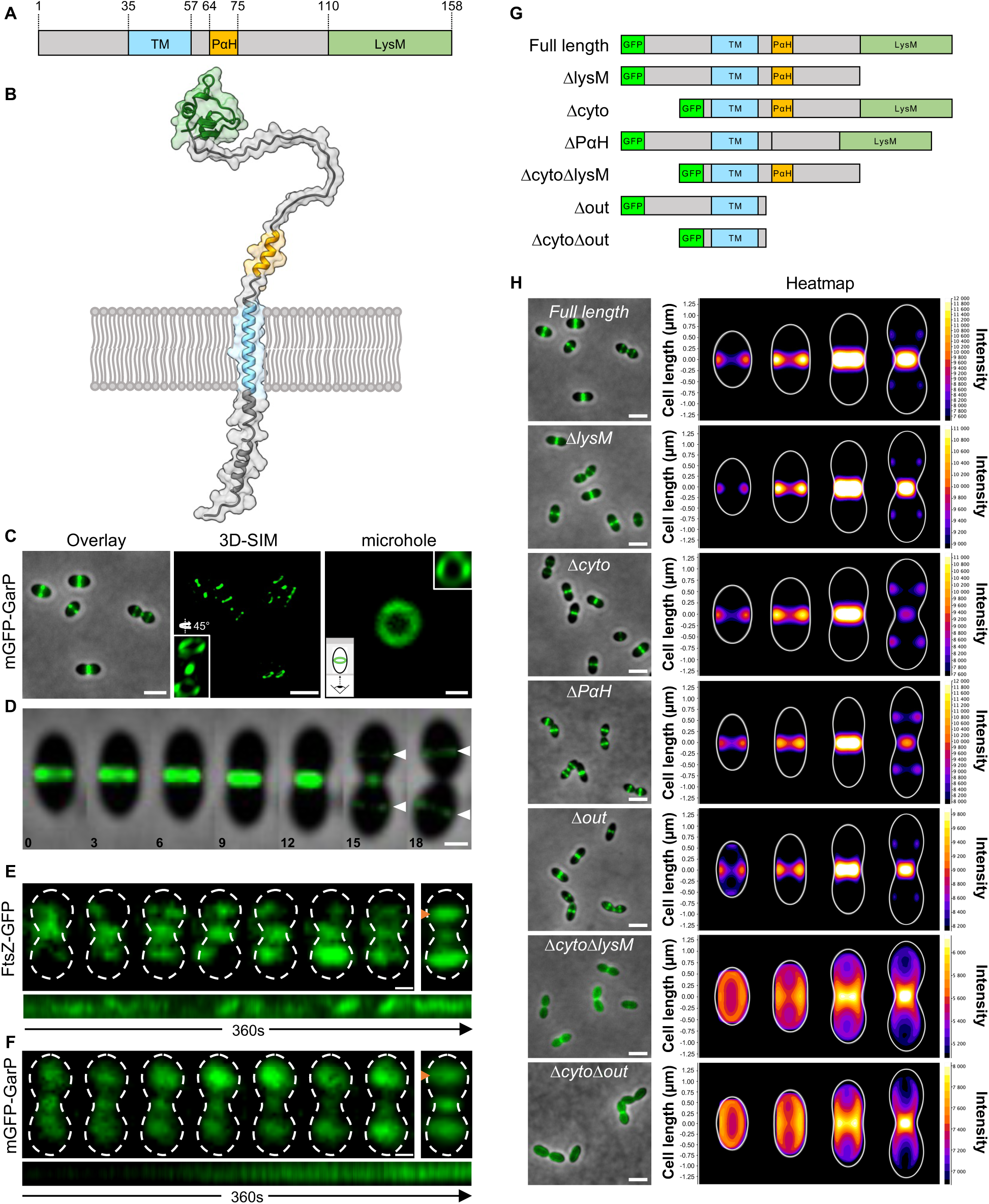
GarP is a membrane protein that forms a ring at midcell. **(A)** Schematic representation and **(B)** Alphafold3 prediction of GarP. The predicted transmembrane (TM) segment and the LysM domain are shown in blue and green, respectively. The predicted α-helix (PαH) is shown in orange. Intrinsically disordered regions are shown in gray. **(C)** Localization of GarP in wild-type cells. Left panel: overlay between phase contrast and GFP fluorescent signal of mGFP- GarP cells. Scale bar, 2 µm. Middle panel: three-dimensional structured illumination microscopy (3D- SIM) snapshot of GarP rings. Scale bar, 2 µm. Right panel: representative conventional and 3D-SIM (inset) images of GarP rings observed in cells trapped in microhole agarose pads. Scale bar, 0.5 µm. **(D)** Representative montage of images acquired by time-lapse microscopy, showing the localization of mGFP-GarP as the cell elongates and divides. Equatorial positions corresponding to the future division sites of the daughter cells are highlighted by a white triangle. Time is indicated in min. Scale bar, 0.5 µm. **(E-F)** Representative montage of images showing the dynamic of FtsZ-GFP (E) or mGFP- GarP (F) at the parental and future division sites of wild-type cells observed by TIRF microscopy for 360 s (9 s intervals). A summation of all the images and a kymograph are shown on the right and below the montage, respectively. The kymograph (1 frame/sec) was generated from a line across one of the equatorial sites, highlighted by an orange triangle on the summation images. Scale bar, 0.5 µm. **(G)** Schematic topology of the full-length mGFP-GarP fusion and derivatives. GFP is indicated by a bright green box. **(H)** Images showing the localization of the mGFP-GarP fusion (Full length) and derivatives. Merged images between the GFP and the phase contrast channels are shown on the left. Scale bars, 2 µm. Corresponding averaged sub-cellular localization of the mGFP-GarP fusion and derivatives are shown as heatmaps on the right. The sub-cellular localization of each variant was monitored for 4 classes of cell length as a proxy for their progression through the cell cycle. The color saturation represents the density of the localizations using a Fire LUT. Localization experiments were performed in triplicate on at least 1000 cells for each fusion.

To study GarP, we first generated an N-terminal GFP fusion (mGFP-GarP) that was stable and fully functional (Extended Data Fig. 1a-d). Using conventional or structured illumination microscopy (SIM), we observed that mGFP-GarP forms a ring at the division site (Fig. 1c). Moreover, mGFP-GarP localizes at future division sites of the daughter cells (called equatorial positions) when the division of the mother cell (the parental division) is almost finished, suggesting that GarP might be a late division protein (Fig. 1d). To gain insights into the dynamics of GarP at the division site, individual *S. pneumoniae* cells expressing mGFP-GarP or FtsZ-GFP were observed using both total internal reflection fluorescence (TIRF) microscopy (Fig. 1e, f) or conventional fluorescence microscopy with cells positioned vertically in agarose microholes (Extended Data Fig. 1e, f). In both cases, mGFP-GarP exhibited no circumferential movement compared to FtsZ-GFP, indicating that GarP does not display the FtsZ treadmilling or bPBPs movement^1^. We next sought to determine which part of GarP is required for its localization to the division site and generated a series of truncated GarP mutants (Fig. 1g). All mutant forms of GarP were stable and expressed as the only source of GarP from the native chromosomal locus (Extended Data Fig. 1g). Surprisingly, single deletion of either the cytoplasmic domain, the entire extracellular domain, the LysM, or the PαH domains did not significantly affect the localization of GarP (Fig. 1h). However, deletion of the cytoplasmic domain in combination with that of the LysM domain or the entire extracellular domain resulted in a significant delocalization throughout the cell periphery, with a remaining enrichment at the septum (Fig. 1h). This indicates that multiple domains contribute to the positioning of GarP at the division site.

### GarP is required for pneumococcal cell shape

To assess the role of GarP in the pneumococcus, we then constructed a Δ*garP* deletion mutant and examined the effect on cell growth and morphology. The Δ*garP* strain was obtained with a normal transformation efficiency, suggesting that *garP* is not essential and that the deletion strain did not acquire suppressor mutations. The deletion of *garP* did not affect the growth rate but caused premature and faster cell lysis (Fig. 2a), suggesting a higher sensitivity to autolytic enzymes^27^. Phase contrast and scanning electron microscopy (SEM) further revealed cells with abnormal shapes and sizes (Fig. 2b). Automated image analysis revealed that Δ*garP* mutant cells are significantly thinner than wild-type cells and that the distribution of the cell length is more dispersed (Fig. 2c). Normal cell shape and growth were restored when Δ*garP* was complemented from an ectopic chromosome locus (Δ*garP*; PcomX-FLAG-*garP*) (Extended Data Fig. 2). Strikingly, the deletion of *garP* was also consistently accompanied by the formation of spherical minicells of varying size (diameter from 0.20 to 0.74 μm, measured on 100 minicells). When Δ*garP* cells expressing a fluorescent membrane probe were observed by SIM, the presence of septa close to the tip of the cell pole (Fig. 2d) further suggested that the number of minicells was likely underestimated. In total, these observations show a clear phenotype for the removal of *garP* and that this protein is required for normal cell morphogenesis and division, as well as resistance toward autolytic enzymes.

**Fig. 2:**
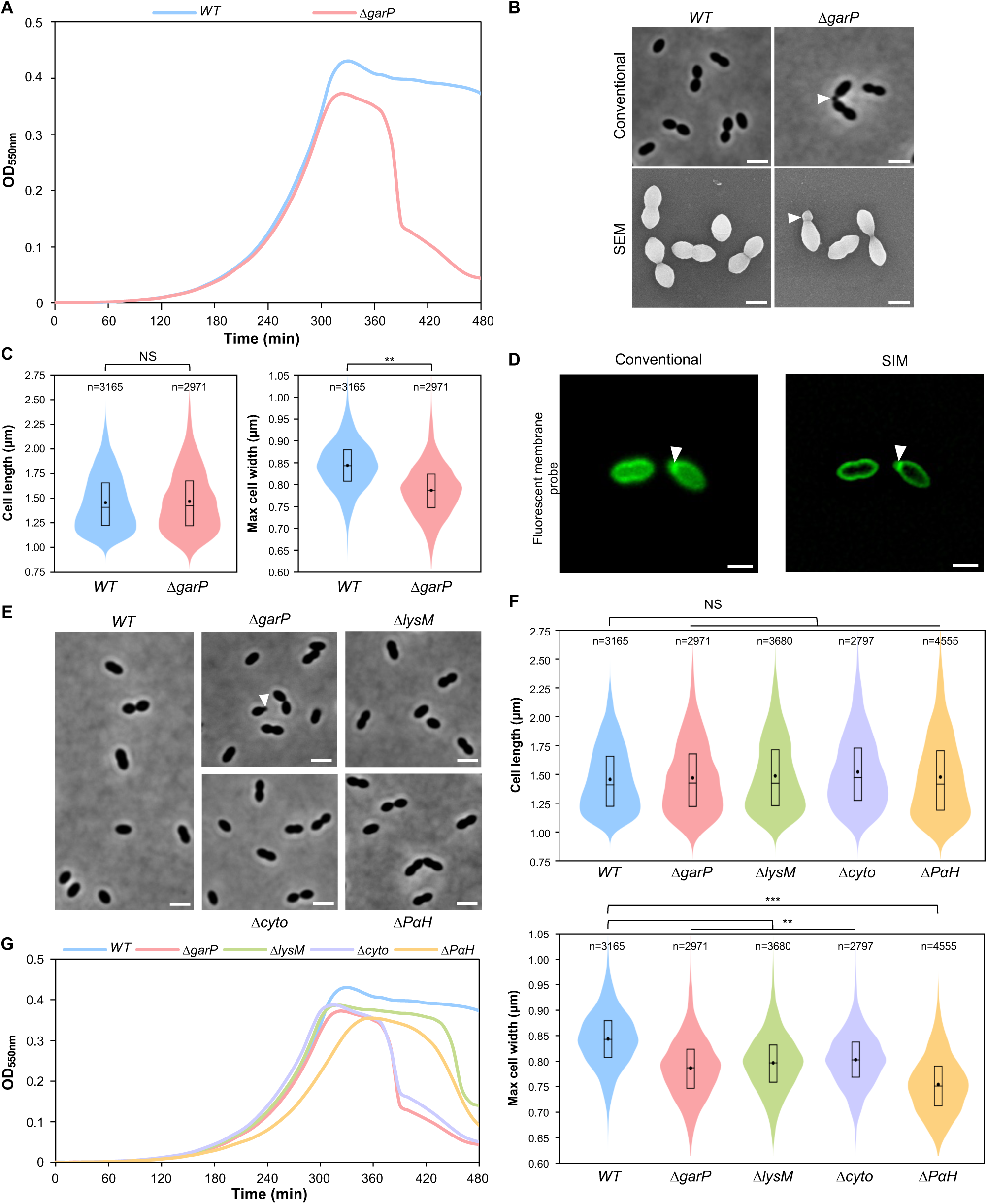
Cell morphogenesis is hindered in the absence of GarP. **(A)** Growth of wild-type (WT) and *ΔgarP* strains. **(B)** Phase contrast (conventional) and electron microscopy (SEM) images of WT and *ΔgarP* cells. Minicells are highlighted by a white triangle. Scale bars, 2 µm (top panels) and 1 µm (lower panels). **(C)** Violin plots showing the distribution of the cell length and maximum cell width from three independent experiments, with n indicating the total number of cells. The box indicates the 25^th^ to the 75^th^ percentile. The mean and the median are indicated with a dot and a line in the box, respectively. Statistical comparison was done using a t-test. **, p <0.01 and NS, not significant, p > 0.05. **(D)** Representative images of *ΔgarP* cells expressing a GFP- tagged membrane probe observed by conventional or structured illumination microscopy (SIM). The white triangle highlights the formation of a septum at the tip of the cell pole. Scale bars, 1 µm. **(E)** Phase contrast microscopy images of WT, Δ*garP* and derivatives (*garP_ΔlysM_*, *garP_Δcyto_*, *garP_ΔPαH_)* strains. Minicells are highlighted by a white triangle. Scale bars, 2 µm. **(F)** Violin plots showing the distribution of the cell length and maximum cell width from three independent experiments, with n indicating the total number of cells. Statistical comparison was done using a t-test. The box indicates the 25^th^ to the 75^th^ percentile. The mean and the median values are indicated with a dot and a line in the box, respectively. ***p < 0.001, **p < 0.01 and NS, not significant, p > 0.05. **(G)** Growth of WT, Δ*garP* and derivatives (*garP_ΔlysM_*, *garP_Δcyto_*, *garP_ΔPαH_)* strains in C+Y medium at 37°C.

To assess the importance of the different domains of GarP, we analyzed the ability of the truncated GarP derivatives to support cell morphology and growth. Cells expressing either GarP_Δcyto_, or GarP_ΔLysM_, or GarP_ΔPαH_ showed the same morphological defects as the Δ*garP* cells, suggesting that they are essential for GarP function (Fig. 2e, f). However, it can be observed that the lysis upon entry into the stationary phase was slower in the absence of the LysM domain or the PαH helix compared to the Δ*garP* mutant (Fig. 2g). Importantly, only the deletion of the PαH helix significantly reduced the growth rate compared to the other *garP* mutants, suggesting that this potential α-helix might play a critical function (Fig. 2g).

### Minicell formation results from aberrant ectopic division septa with reduced nascent PG

To examine the formation of minicells, we looked at Δ*garP* cells at the single-cell level using time- lapse microscopy (Fig. 3a). We observed that minicells always formed from an ectopic and misplaced septum located between the parental division site and the equatorial sites of the daughter cells, resulting in an asymmetrical division (Fig. 3a). To understand the origin of this ectopic positioning of the division site, we examined the subcellular localization and the dynamics of key division proteins. For that, we looked at the localization of MapZ, a protein that positions the division site and recruits FtsZ^28,29^. In a wild-type strain, GFP-MapZ forms a ring at the equator of newborn cells^28^ (Fig. 3b). As the cell elongates, the GFP-MapZ ring splits into two rings that move away from each other at the rate of cell elongation, migrating with the two new cell equators that will eventually become the division site of the two future daughter cells^28^ (Fig. 3b). In the absence of *garP*, MapZ positions properly at parental and equatorial sites. However, we also observed the formation of an additional MapZ ring at the ectopic position that will eventually become the division site generating the minicell (Fig. 3b, c). Note however that the ectopic division plane remains perpendicular to the longitudinal axis of the cell, suggesting that only its longitudinal placement is affected by the deletion of *garP.* We then looked at the localization of FtsZ and two bPBPs (PBP2x, as a proxy for the divisome, and PBP2b as a proxy for the elongasome) in Δ*garP* cells. As shown in Fig. 3b, the 3 proteins localize to the parental division site and the cell equators, but also to the ectopic division site. These experiments show that the deletion of *garP* leads to the recruitment of MapZ, FtsZ, the majors PBPs and likely the entire division and elongation machineries to an ectopic position located between the division site of the mother cell and the future division site of one daughter cell.

**Fig. 3:**
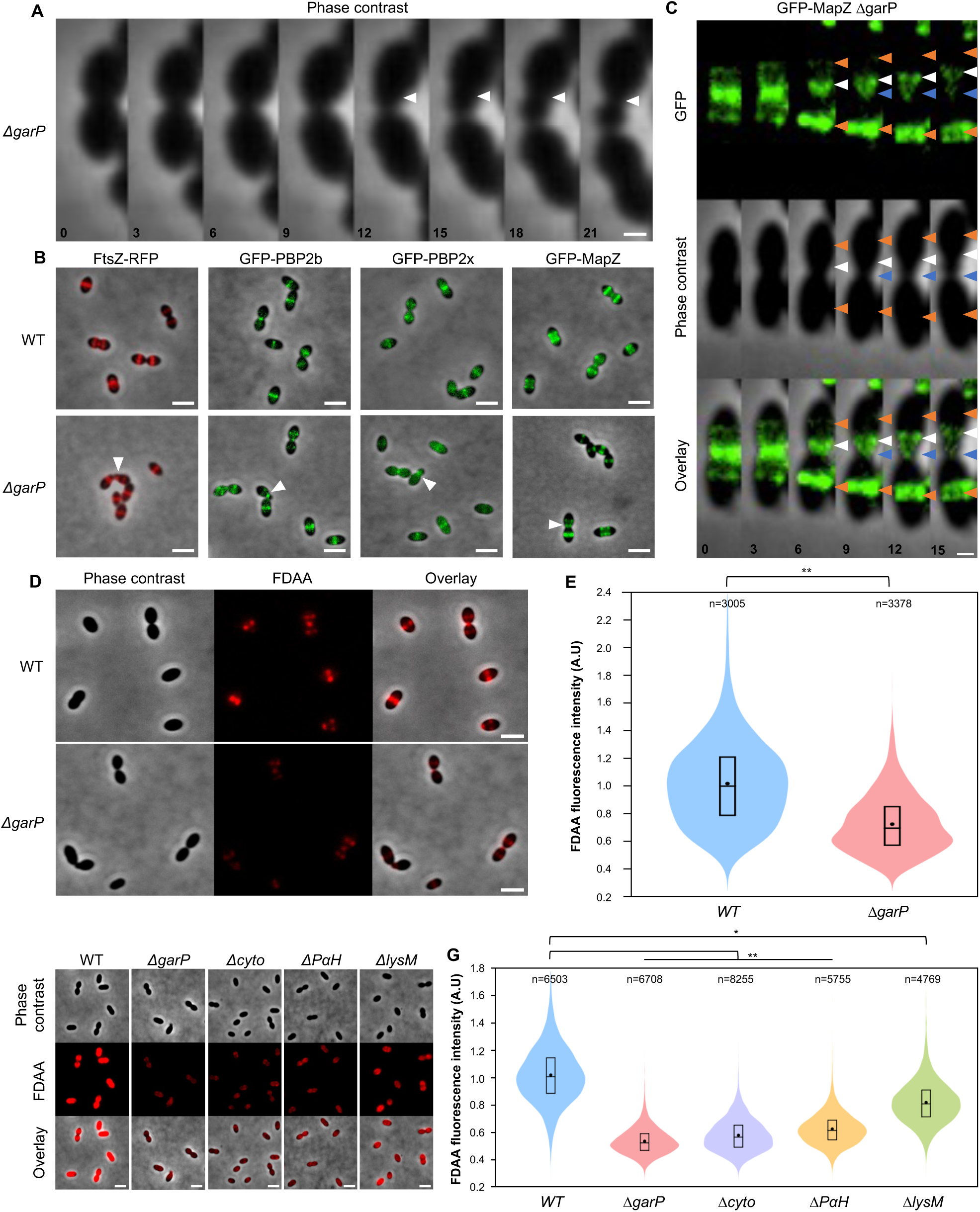
Minicell formation results from aberrant ectopic division septa and the absence of GarP leads to reduced FDAA incorporation. **(A)** Representative montage of images acquired by time-lapse microscopy, showing the formation of a minicell in a *ΔgarP* cell. The position of the ectopic division site is indicated by a white triangle. Time is given in min. **(B)** Images of wild-type (WT) or *ΔgarP* cells expressing the FtsZ-RFP, GFP-PBP2b, GFP-PBP2x or GFP-MapZ fusion. Merged images between the phase contrast and GFP or RFP channels are shown. Ectopic division sites are indicated by white triangles. Scale bars, 2 µm. **(C)** Representative montage of images acquired by time-lapse microscopy, showing the localization of GFP-MapZ as a *ΔgarP* cell elongates. The formation of an ectopic MapZ ring is highlighted by a white triangle. The parental division site and cell equators are highlighted by blue and orange triangles, respectively. Time is given in min. **(D)** Conventional fluorescence microscopy images of wild-type and *ΔgarP* cells after a short labelling pulse with the fluorescent D-amino acid (FDAA) TDL. The phase contrast and the RFP channels, as well as a merged image are shown. Scale bars, 2 µm. **(E)** Violin plots showing the distribution of FDAA fluorescence intensity for wild-type and *ΔgarP* cells after a short pulse of FDAA (TDL) labelling from three independent experiments, with n indicating the total number of cells. The box indicates the 25^th^ to the 75^th^ percentile, and the whiskers indicate the minimum and the maximum values. The mean and the median are indicated with a dot and a line in the box, respectively. Statistical comparison was done using a t-test. **p < 0.01. **(F)** Representative images of wild-type, *ΔgarP* and derivatives (*garP_Δcyto_, garP_ΔPαH,_ garP_ΔlysM_*) cells after a long period of labeling with FDAA (TDL). The phase contrast, the RFP and the merged image between the two channels are shown. Scale bars, 2 µm. **(G)** Violin plots showing the distribution of the FDAA fluorescence intensity for wild-type, *ΔgarP* and derivatives (*garP_ΔlysM_, garP_Δcyto_, garP_ΔPαH_*) cells after a long period of FDAA (TDL) labelling from three independent experiments, with n indicating the total number of cells. The box indicates the 25^th^ to the 75^th^ percentile, and the whiskers indicate the minimum and the maximum values. The mean and the median are indicated with a dot and a line in the box, respectively. Statistical comparison was done using a t-test. **p < 0.01 and *p < 0.05.

MapZ localizes at the cell equators upon interaction with a yet uncharacterized PG motif^30^. We hypothesized that the misplacement of the MapZ ring in the Δ*garP* mutant might be due to a defect in the assembly of the PG layer. To test this hypothesis, we pulse-labeled wild-type and Δ*garP* mutant cells with the fluorescent D-amino acid TAMRA-D-lysine (TDL) probe, which is incorporated into the PG by the transpeptidase activity of the PBPs^31^, and tracked its incorporation during cell division (Fig. 3d). In both strains, TDL incorporation was observed at parental and equatorial positions but surprisingly, the intensity of the signal was significantly lower in Δ*garP* cells (Fig. 3e). Compared to the wild-type strain, a weaker fluorescent signal was also observed when Δ*garP* cells were labeled for a longer period of time (Extended Data Fig. 3), indicating that the rate of incorporation of TDL is affected by the absence of GarP. We also evaluated the contribution of the cytoplasmic, the LysM, and the PαH domains to PG labeling (Fig. 3f, g). While the deletion of the LysM domain had a modest impact on TDL incorporation, the deletion of either the cytoplasmic or the PαH helix strongly decreased the fluorescence intensity. These results therefore suggest that GarP is involved in the regulation of PG synthesis in *S. pneumoniae*. Notably, the cytoplasmic and the PαH helix of GarP are essential for this function.

### GarP interacts with PBP1a and promotes its activity

FDAA are incorporated by the PBPs, through the exchange of the terminal D-Ala residue carried by the pentapeptides of the PG^31^. To understand how FDAA incorporation is affected in the absence of *garP*, we aimed to identify the PBP(s) responsible for TDL incorporation in *S. pneumoniae*. Since the deletion of *garP* did not lead to cell elongation or constriction defects (Fig. 2b, c), it is unlikely that a SEDS-bPBP activity is directly affected^11,32^. Therefore, we pulse-labeled non-essential aPBP mutants (Δ*pbp1a*, Δ*pbp2a*, Δ*pbp1b*) with TDL and tracked its incorporation during cell division. Strikingly, only the deletion of *pbp1a* caused a significant reduction in TDL incorporation, reaching levels similar to those observed with the *garP* mutant (Fig. 4a, b). The same observation was made with long periods of TDL labeling (Extended Data Fig. 4a, b), suggesting that GarP may stimulate PBP1a activity *in vivo*.

**Fig. 4:**
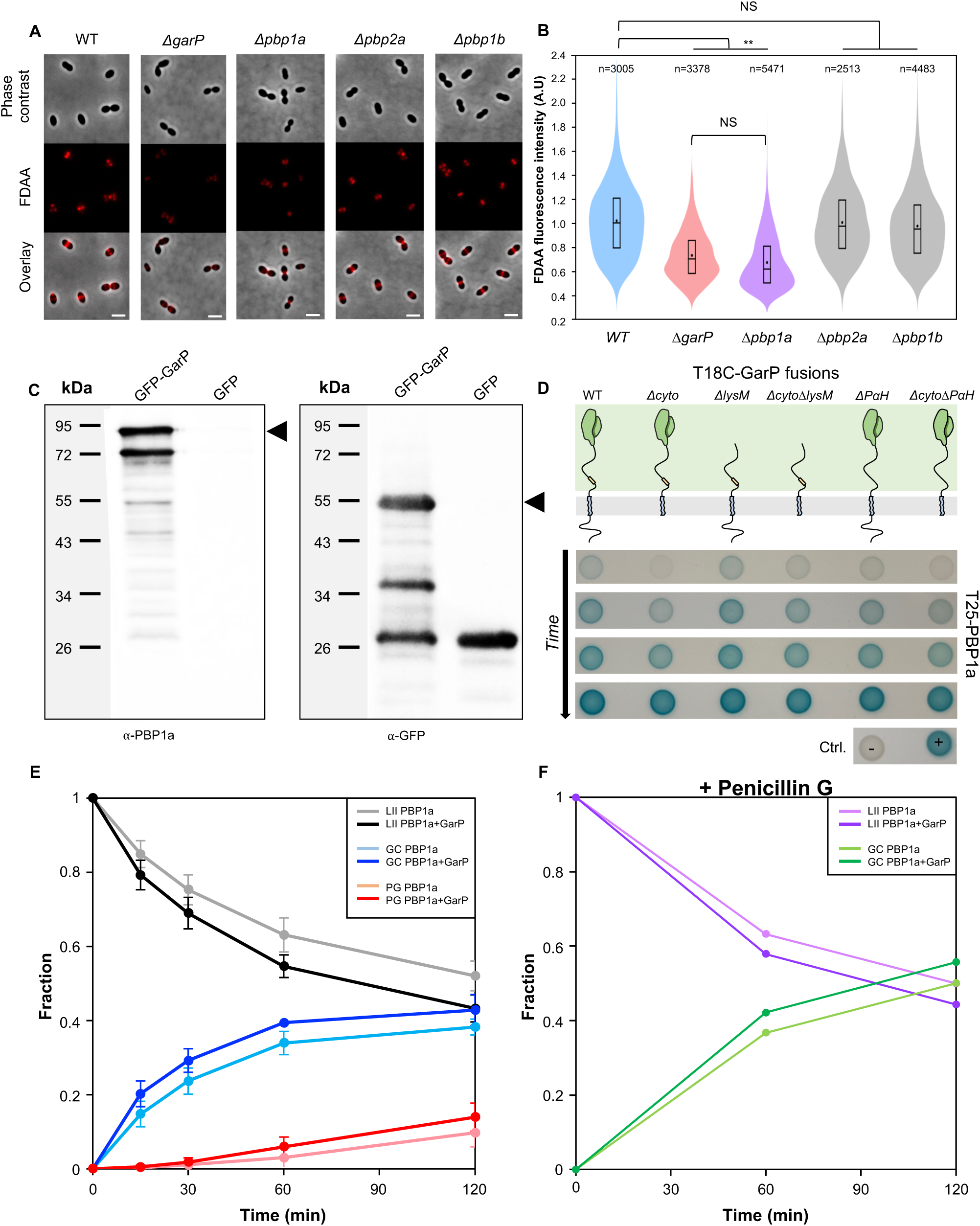
GarP interacts with and promotes the activity of PBP1a. **(A)** Representative images of wild-type, *ΔgarP, Δpbp1a*, *Δpbp2a* and *Δpbp1b* cells after a short pulse of FDAA (TDL) labelling. The phase contrast, the RFP and the merged images between the RFP and the phase contrast channels are shown. Scale bars, 2 µm. **(B)** Violin plots showing the distribution of the normalized FDAA fluorescence intensity for wild-type, *ΔgarP, Δpbp1a*, *Δpbp2a* and *Δpbp1b* cells after a short pulse of FDAA labelling, from three independent experiments, with n indicating the total number of cells. The box indicates the 25^th^ to the 75^th^ percentile. The mean and the median are indicated with a dot and a line in the box, respectively. Statistical comparison was done using a t-test. **p < 0.01 and ns, not significant, p > 0.05. **(C)** Co-immunoprecipitation of PBP1a with mGFP-GarP and GFP using anti-GFP antibodies. Samples were analyzed by immunoblotting using anti-PBP1a antibodies (left panel) to determine the presence of co-immunoprecipitated PBP1a or anti-GFP antibodies (right panel) to check the specificity of the anti-GFP immunoprecipitation. The data shown are representatives of experiments made independently in triplicate. **(D)** Bacterial two-hybrid analyses. Plasmids, expressing the T25 fragment of the adenylate cyclase protein fused to the N-terminus of PBP1a or the T18 fused to the N-terminus of GarP and its derivatives (GarP_ΔlysM_, GarP_Δcyto_, GarP_ΔlysMΔcyto_, GarP_ΔPαH_, GarP_ΔPαHΔcyto_), were co-transformed in *E. coli* BTH101. The blue coloration indicates positive interactions. The colonies were pictured every 5 h after 24 h of incubation at 25°C. The data shown are representatives of experiments made independently in triplicate. **(E)** Time course of peptidoglycan assembly by PBP1a in presence or absence of GarP. The reaction was analyzed by SDS-PAGE and the gel was then imaged under UV-transillumination at various time intervals. The fluorescence intensity in the different areas of the gel (Extended Data Fig. 4d) corresponding to the Lipid II (LII), uncross-linked glycan chains (GC) and cross-linked peptidoglycan (PG) was quantified and represented as fraction of the total fluorescence for each sample. LII is in grey or black, GC in lighter blue or deeper blue, PG in salmon or red in presence or absence of GarP, respectively. The average of three independent experiments is shown with the standard error of the mean. **(F)** Same experiment as described in (G), performed in presence of penicillin G to block TP activity. LII is in light or deep purple, GC in light or deep green in presence or absence of GarP, respectively.

To investigate this further, we conducted three complementary approaches. As previously demonstrated, *pbp1a* exhibits synthetic lethality with *pbp2a*, *macP*, and *gpsB*, and has a synthetic viable relationship with *mpgA*^23,33^. We reasoned that if GarP is required for PBP1a activity, they should share the same genetic interactions. Consistent with this, *garP* also shows synthetic lethality with *pbp2a*, *macP*, and *gpsB*, and shares the synthetic viable relationship with *mpgA* (Extended Table 1), demonstrating a strong functional link between *pbp1a* and *garP*. Next, we performed co- immunoprecipitations using detergent-solubilized membrane preparations from *S. pneumoniae* cells to determine whether GarP interacts with PBP1a. We immunoprecipitated mGFP-GarP using anti-GFP antibody resin and revealed the presence of PBP1a using a specific antibody. PBP1a efficiently co- precipitated with mGFP-GarP, but not in control preparations derived from cells expressing only the mGFP (Fig. 4c). To determine if GarP directly interacts with PBP1a, we used an *E. coli* bacterial two- hybrid system to screen for interactions between PBP1a and different versions of GarP. In agreement with co-immunoprecipitations, a strong interaction between PBP1a and GarP was detected (Fig. 4d). Interestingly, this interaction was not affected by the deletion of the LysM domain, and only partially affected upon single deletion of the cytoplasmic domain or the PαH helix, suggesting some redundancy in their ability to mediate the interaction between PBP1a and GarP. This interaction was further affected, but not completely abolished, upon the double deletion of the cytoplasmic domain and the PαH helix (Fig. 4d). Consequently, we cannot exclude a significant role for the TM segment of GarP in the interaction with PBP1a. We conclude that GarP forms a stable complex with PBP1a through cumulative interactions involving the TM segment, the cytoplasmic and the PαH domains.

Finally, we assessed the biochemical activity of PBP1a in presence and in absence of GarP. For that, we first purified PBP1a^34^ and GarP independently (Extended Data Fig. 4c). Then, we incubated recombinant PBP1a with a mixture of unlabeled lipid II and fluorescent lipid II in presence and in absence of GarP^34^. A modest but reproducible stimulation of PG assembly by PBP1a was observed in the presence of GarP (Fig. 4e and Extended Data Fig. 4d). In addition, the faster consumption of lipid II and appearance of glycan chains suggested that GarP stimulates mostly the GT activity of PBP1a. A time course reaction performed in the presence of penicillin G, which inhibits the TP activity, confirmed that the GT activity of PBP1a is enhanced by GarP (Fig. 4f). It is thus likely that the enhanced PG assembly observed in the absence of penicillin and in the presence of GarP results from the stimulated GT activity. Collectively, all of these experiments demonstrate that GarP promotes the activity of PBP1a through a direct interaction.

### The PαH helix of GarP is required for PBP1a activation

To understand how GarP can stimulate the activity of PBP1a at the molecular level, we used AlphaFold3 to predict the GarP-PBP1a complex (Fig. 5a-d). Although the location of the LysM domain varied widely depending on the prediction model, AF3 consistently predicted the PαH helix interacting with a surface-exposed groove on the GT domain of PBP1a. In this model, the PαH helix of GarP is adjacent to the helix of PBP1a containing the E91 catalytic residue, according to the structure of PBP1b of *Escherichia coli*^35^, with potential interactions between several amino acids (Fig. 5a-d and Extended Data Fig.5). Together with our observations regarding the importance of the PαH in TDL incorporation and growth rate (Fig. 2g and Fig. 3f, g), we hypothesized that the PαH may be critical for stimulating PBP1a activity. To test this, we individually mutated each residue of the PαH domain (64-75) to alanine and evaluated their effect on pneumococcal cell morphology (Fig. 5e, f). Only the L68A and F71A mutations induced morphological and growth defects similar to that of the Δ*PαH* mutant. (Fig. 5g). Last, we analyzed L68A and F71A mutants for their ability to incorporate TDL. As shown in Fig. 5h, a significant reduction comparable to that of the PαH deletion was detected. These observations indicate that the PαH domain, and particularly L68 and F71, are crucial for the stimulation of PBP1a by GarP (Fig. 5d). Supporting this, single or double mutations of L68A and F71A had the same synthetic lethality with *pbp2a* as a g*arP* deletion, although some clones appeared after 48 h of incubation *(see below)* (Extended Table 2). These observations are thus consistent with an activation of PBP1a by the PαH helix of GarP.

**Fig. 5:**
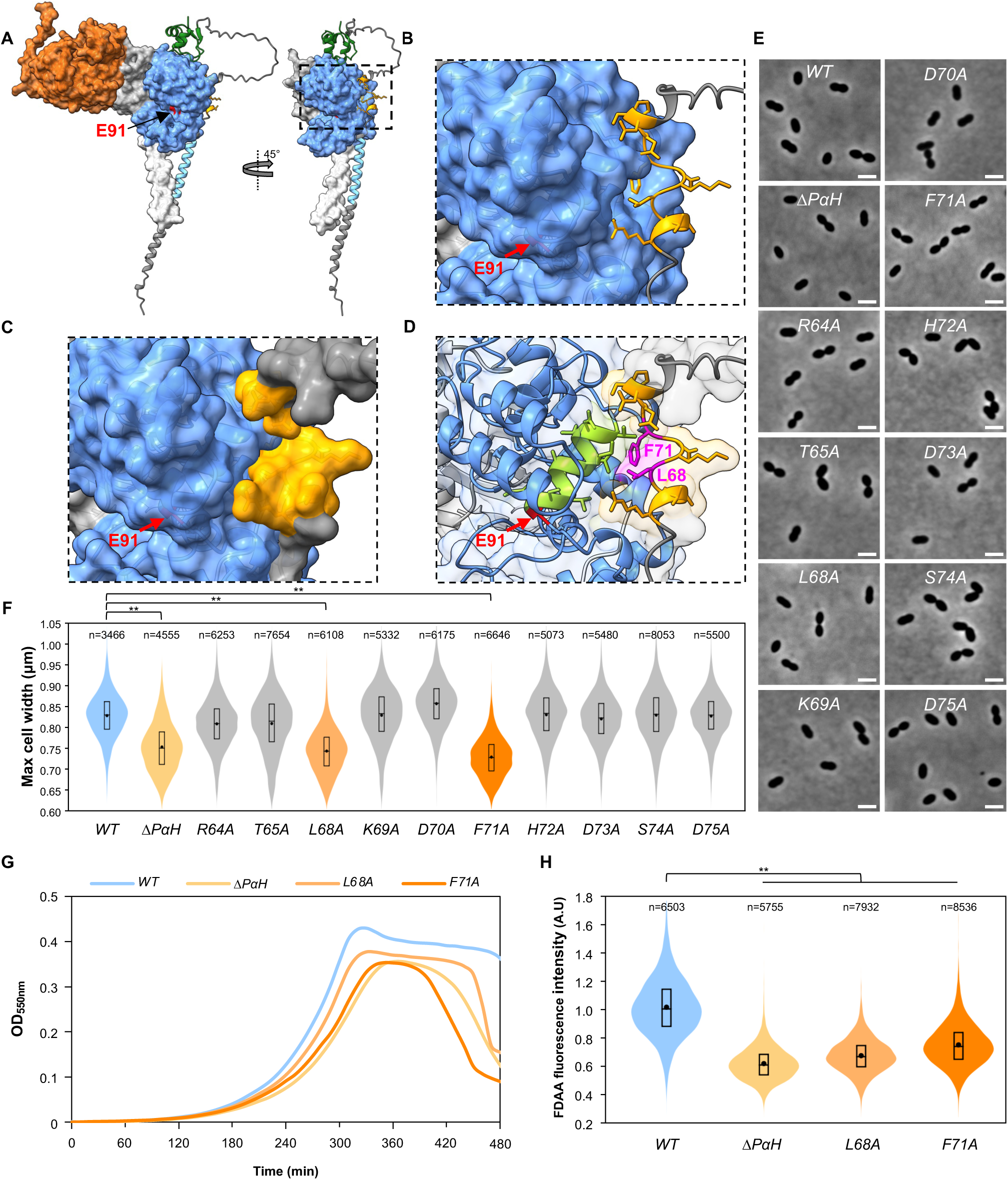
The PαH of GarP is required for PBP1a activation. **(A)** The complex between PBP1a and GarP predicted by Alphafold3. PBP1a is represented with a semi-transparent surface while GarP is represented as a ribbon diagram. The N-terminal cytoplasmic region and the TM segment of PBP1a are shown in white. The GT and TP domains of PBP1a are shown in blue and orange, respectively, separated by the linker region shown in light gray. The flexible C-terminal domain (S669-P719) is not shown for better clarity. The catalytic residue (E91) of PBP1a GT domain is highlighted in red. The predicted transmembrane segment and the LysM domain of GarP are shown in blue and green, respectively. The predicted α-helix (PαH) of GarP is shown in light orange. Others regions (intrinsically disordered) of GarP are shown in gray. **(B) (C)** Zoom-in on the region boxed in (A), highlighting the interaction surface between the PαH of GarP and the GT domain of PBP1a. **(D)** Zoom-in on the predicted interaction between the PαH of GarP and the helix of GT domain of PBP1a bearing the catalytic residue (E91) highlighted in red. The GT domain of PBP1a is shown in blue and the helix bearing the catalytic residue is shown in light green. The predicted α-helix of GarP is shown in light orange and the two residues important for PBP1a stimulation (L68 and F71) are shown in purple. **(E)** Phase contrast microscopy images of wild-type (WT) and PαH single mutation strains. Scale bars, 2 µm. **(F)** Violin plots showing the distribution of the maximum cell width of WT and PαH single mutation strains, from three independent experiments, with n indicating the total number of cells. The box indicates the 25^th^ to the 75^th^ percentile. The mean and the median are indicated with a dot and a line in the box, respectively. Statistical comparison was done using a t-test. **p <0.01. **(G)** Growth of WT, *GarP-ΔPαH*, L68A and F71A mutant strains. **(H)** Violin plots showing the distribution of the FDAA fluorescence intensity for wild-type, *garP_ΔPαH_*, *garP*_L68A_ and *garP*_F71A_ cells after a long period of FDAA labelling, from three independent experiments, with n indicating the total number of cells. The box indicates the 25^th^ to the 75^th^ percentile. The mean and the median are indicated with a dot and a line in the box, respectively. Statistical comparison was done using t-test. **p < 0.01.

### The PBP1a-A124T mutation compensates for the lack of GarP

As mentioned above, the deletion of *pbp2a* in the GarP_L68A-F71A_ background resulted in the apparition of clones after 48 h of incubation, indicative of suppressive mutations. Since *pbp2a* deletion is synthetically lethal with Δ*pbp1a*^22^, we reasoned that these suppressive mutations might enhance PBP1a activity, by-passing the need for GarP. To identify these compensatory mutations, we sequenced both the *garP* locus, to exclude those with reverse mutations in *garP,* and the *pbp1a* locus. Strikingly, we found a single point mutation (A124T) in the GT domain of PBP1a. To determine if the A124T point mutation can bypass the need for GarP, this allele was backcrossed into the parental strain or the *ΔgarP* mutant and their genetic interaction with *pbp2a* was tested. As expected, deletion of *pbp2a* was readily obtained in the *ΔgarP pbp1a*-A124T strain but not in the *ΔgarP* strain (Extended Table 3). We also examined the effect of the *pbp1a*-A124T allele on bacterial growth, cell morphology and TDL incorporation (Fig. 6a-d). The cells harboring the *pbp1a*-A124T allele showed a slightly better growth rate and a reduced propensity for autolysis upon entry into the stationary phase compared to the wild-type cells (Fig. 6c). Accordingly, the Δ*garP* mutant harboring the *pbp1a*-A124T allele also showed a better growth rate and a higher maximum optical density compared to the *ΔgarP* mutant. The stationary phase remained however comparable to that of the Δ*garP* mutant, indicating that the A124T allele does not fully compensate for the absence of GarP. Importantly, introducing the *pbp1a*- A124T allele did not cause any morphological defects in wild-type or Δ*garP* cells (Fig. 6a, b). Moreover, while cells expressing the *pbp1a*-A124T allele exhibited TDL incorporation similar to the WT strain, *ΔgarP* cells expressing the *pbp1a*-A124T allele showed improved TDL incorporation compared to *ΔgarP* cells (Fig. 6d). These observations strongly suggest that the A124T mutation stimulates PBP1a activity and can bypass, at least partially, the requirement for GarP.

**Fig. 6:**
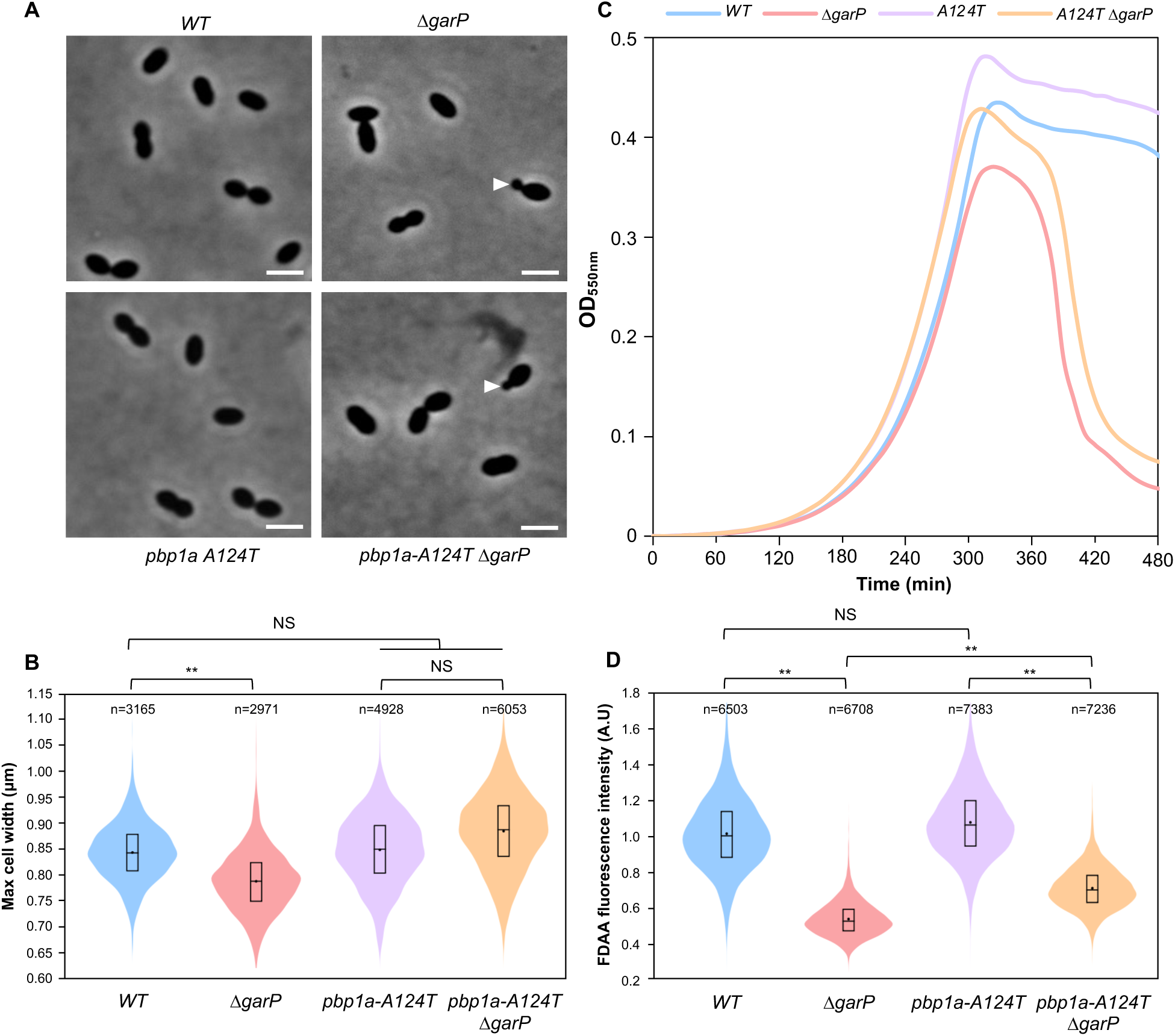
The PBP1a-A124T mutation by-passes the requirement of GarP. **(A)** Phase contrast microscopy images of wild-type and *pbp1a_A124T_* cells in presence or absence of *garP*. Minicell formation is indicated by white triangles. Scale bars, 2 µm. **(B)** Violin plots showing the distribution of the maximum cell width from three independent experiments, with n indicating the total number of cells. The box indicates the 25^th^ to the 75^th^ percentile. The mean and the median are indicated with a dot and a line in the box, respectively. Statistical comparison was done using a t-test. **p < 0.01 and NS, not significant, p > 0.05. **(C)** Growth of wild-type, *ΔgarP*, *pbp1a_A124T_* and *pbp1a_A124T_ ΔgarP* mutants in C+Y media at 37°C. **(D)** Violin plots showing the distribution of the FDAA fluorescence intensity for wild-type, *ΔgarP*, *pbp1a_A124T_* and *pbp1a_A124T_ ΔgarP* mutants cells after a long period of FDAA labelling, from three independent experiments, with n indicating the total number of cells. The box indicates the 25^th^ to the 75^th^ percentile. The mean and the median are indicated with a dot and a line in the box, respectively. Statistical comparison was done using a t-test. **p < 0.01 and NS, not significant, p > 0.05.

### The PBP2a GpsB-binding motif SRxxR is conserved in GarP, MpgA and PgdA

As mentioned above, PBP2a interacts with GpsB through its [(S/T)RxxR(R/K)] motif^17^. Remarkably, we found that the cytoplasmic domain of GarP also contains the consensus motif [(S/T)RxxR(R/K)] with the sequence SRAGRR, suggesting that GarP may also interact with GpsB (Fig. 7a). We thus tested the interactions between GpsB and GarP by bacterial two-hybrid (Fig. 7b). Our analysis revealed an interaction between GarP and GpsB, which was abolished when the SRAGRR sequence was deleted. Interestingly, we identified two other proteins with a conserved (S/T)RxxR(R/K) motif : the PG deacetylase PgdA (SRLGRG) and the muramidase MpgA (SRRSERR) (Fig. 7a). We then determined whether these proteins interact with GpsB and whether the interaction is mediated by their SRxxR(R/K) motif, using wild-type or truncated versions of PgdA and MpgA (Fig. 7b). As expected, our analysis revealed strong interactions between GpsB and PgdA or MpgA that were abolished when their respective (S/T)RxxR(R/K) motifs were deleted.

**Fig. 7:**
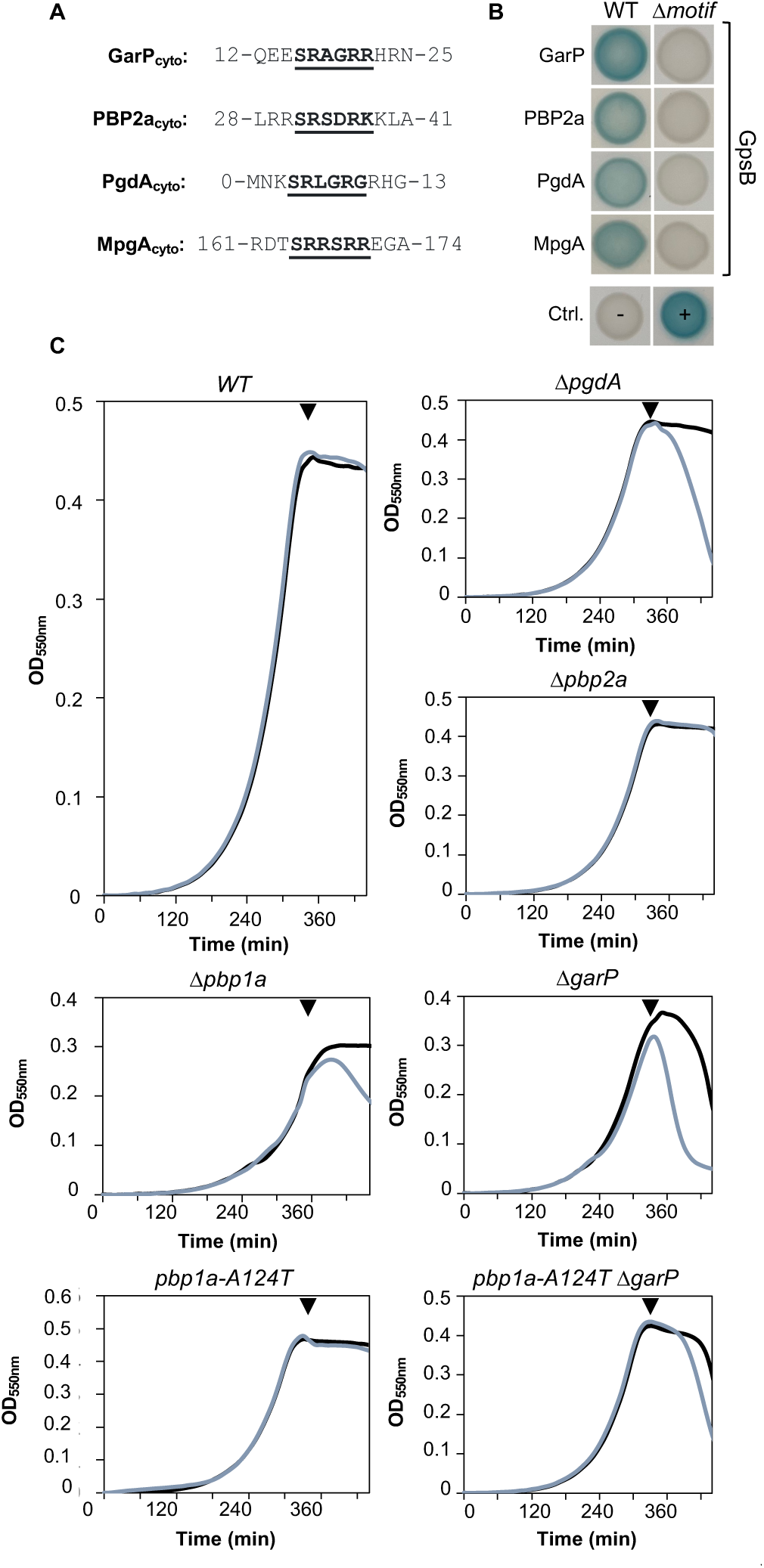
A GpsB-binding motif SRxxR is conserved in GarP, PBP2a, PgdA and MpgA. **(A)** Local alignment of the amino acid sequence of the SRxxR motifs observed in the cytoplasmic domain of GarP, PBP2a, PgdA and MpgA. The motif is highlighted in bold and underlined. **(B)** Bacterial two-hybrid analyses. Plasmids expressing the T25/ T18 fragment of the adenylate cyclase protein fused to the N-terminus or the C-terminus of GpsB, or the T25/T18 fragment fused to the N-terminus of GarP, PBP2a, PgdA or MpgA and their derivatives in which the SRxxR motif was deleted, were co-transformed in *E. coli* BTH101. The picture was taken after 48 h of incubation at 25°C. The blue coloration indicates positive interactions. The data shown are representatives of experiments made independently in triplicate. **(C)** Lysozyme sensitivity of wild-type*, ΔpgdA, ΔgarP,Δpbp1a, Δpbp2a, pbp1a_A124T_* and *pbp1a_A124T_ ΔgarP* strains. The growth in C+Y media at 37°C was monitored with or without the addition of chicken egg lysozyme (100 µg/mL) at the onset of the stationary phase (black triangle).

PgdA is an N-acetylglucosamine deacetylase that makes PG resistant to the hydrolytic action of lysozyme^36^. The latter is abundant in the mucosal surface of upper respiratory tract and PgdA is thus considered as a virulence factor required for host colonization^37^. Since PgdA interacts with GpsB (Fig. 7b) and might be part of a GarP/PBP1a/PBP2a/MpgA complex, we investigated the role of these other GpsB partners in the sensitivity to lysozyme. To test this, wild-type and mutant strains were grown, and lysozyme was added in the culture at the onset of the stationary phase (Fig. 7c). Strikingly, wild-type and *Δpbp2a* mutant strains were not affected by the addition of lysozyme, but the *ΔgarP* and the *Δpbp1a* mutant strains began to lyse rapidly upon addition of lysozyme, suggesting that N-acetylation might be affected in the absence of GarP or PBP1a. To further investigate the influence of GarP and PBP1a on the sensitivity to lysozyme, we assessed the sensitivity of the *ΔgarP* strain expressing the *pbp1a-A124T* allele that bypasses the need for GarP. This strain showed an intermediate level of lysozyme sensitivity (Fig. 7c). Altogether, this shows that the sensitivity of *S. pneumoniae* to lysozyme is not only affected by PgdA, but also by PBP1a activity, suggesting a functional link between PgdA and PBP1a activities.

MpgA is a muramidase proposed to release PG strands synthesized by PBP1a for cross-linking by the RodA-PBP2b complex^33^. As previously mentioned, *mpgA* has a synthetic viable genetic interaction with *pbp1a* and *garP*, but not with *pbp2a,* suggesting that the deletion of *mpgA* is possible only when PBP1a is absent or inactive (Fig. 4b). Since we demonstrated that lysozyme sensitivity is strongly affected when PgdA or PBP1a is absent (Fig. 7c), we reasoned that *pgdA* could also have a synthetic viable genetic interaction with *mpgA*. In agreement with this hypothesis, we were able to readily delete *mpgA* in the Δ*pgdA* strain (Extended Table 4). Altogether, this suggests that MpgA activity might be dispensable when cells produce fully N-acetylated glycans, but might be required to hydrolyze N-deacetylated glycans that may become toxic if too abundant. It further implies that MpgA activity might require a tight coordination with PBP1a and/or PgdA activity. This finally suggests that GarP, PBP1a, PgdA, MpgA, PBP2a and its cognate regulator MacP, could be part of a complex coordinated by the scaffolding protein GpsB and required for PG remodeling and maturation.

## DISCUSSION

aPBPs play a critical role in the remodeling of the PG layer and/or in repairing damaged areas, ensuring that the PG matrix remains intact and maintains the cell shape^38^. To remodel the PG, aPBPs likely work in concert with PG hydrolases and other modifying enzymes^39^. In *S. pneumoniae*, the two major aPBPs, PBP1a and PBP2a, form a synthetically lethal pair, indicating that they fulfill both specific and overlapping functions. It is therefore essential that the activity of these two aPBPs is regulated and coordinated in order to ensure that the PG layer is correctly remodeled or repaired. Nevertheless, how PBP1a and PBP2a coordinate with each other, with hydrolases and other PG modifying enzymes, has remained completely unknown. The identification and characterization of GarP not only unveils a regulation mechanism for PBP1a, but also, and most importantly, identifies a GpsB-interacting collection of proteins composed of PBP1a and its cognate regulator GarP, PBP2a and its cognate regulator MacP, the PG hydrolase MpgA and the PG deacetylase PgdA. This ensemble of proteins interacts directly (PBP2a, GarP, MpgA and PgdA) or indirectly (PBP1a and MacP) with the hexameric cell division GpsB, either as a single complex or a collection of complexes of varying composition. These or these dynamic complexe(s) would likely control PG remodeling and/or repair during pneumococcal cell elongation and division.

Regarding PBP1a regulation, we demonstrated that GarP directly interacts with PBP1a and promotes its GT activity. We showed that PBP1a activation results from a direct interaction between a predicted α-helix (PαH) of GarP and the GT domain of PBP1a (Fig. 4 and 5). In this heterodimer, the PαH is predicted to interact with a surface-exposed cleft located at the back of the GT domain (Fig. 5a). More precisely, PαH is adjacent to the helix of PBP1a containing the catalytic residue E91, according to AF3 prediction. This interaction surface differs from those reported for other aPBPs with their cognate activators. This is particularly the case for the interaction of *E. coli* PBP1a and PBP1b with the lipoproteins LpoA and LpoB, respectively^13^. Although the two lipoproteins are unrelated in sequence, they both possess a specific C-terminal domain that interacts with their respective aPBP via a non- catalytic domain located in their extracellular region^15^. These non-catalytic domains, designated ODD in PBP1a and UB2H in PBP1b, are also structurally distinct and conserved exclusively in enterobacteria and γ-proteobacteria, respectively^13^. It is proposed that the binding of LpoA to PBP1a- ODD and LpoB-UB2H to PBP1b induces specific conformational changes in the interdomain region between the GT and TP domains of PBP1a and PBP1b, thereby activating their PG polymerization and cross-linking activities^40,41^. A comparable activation mode has been proposed for PBP1b from *Pseudomonas aeruginosa*, which also possesses an U2BH domain, by the lipoprotein LpoP, which is unrelated to the *E. coli* LpoB^42^.

PBP1a from *S. pneumoniae* lacks additional non-catalytic domains such as ODD and UB2H. Instead, its GT domain interacts directly with GarP. How, then, might GarP enhance PBP1a activity? Since L68 and F71 of PαH are crucial for triggering PBP1a activity, it can be hypothesized that they contribute to the optimal orientation and/or stabilization of the PBP1a α-helix carrying the catalytic E91 of the GT domain to promote its activity. A second hypothesis can be formulated by comparing the AF3 models of PBP1a alone and in complex with GarP. A subtle conformational change in the interdomain region of PBP1a is observed upon the binding of GarP, suggesting that the PαH of GarP may also reorient or enhance the movement between the GT and TP domains of PBP1a (Extended Data Fig. 6 and Supplementary Video 1). Last, a third hypothesis concerns a potential structural rearrangement in the GT domain itself. The GT domain of aPBPs is comprised of two lobes, designated the head (upper lobe) and the jaw (lower lobe)^43^. A124 is located in the jaw of the GT domain of PBP1a^44^. In several aPBPs, including *S. aureus* PBP2, the jaw subdomain exhibits conformational flexibility, adopting different conformations when bound to the substrate-mimicking antibiotic moenomycin. This suggests that these structural changes may regulate the enzymatic activity^43^. In *S. pneumoniae,* it has been proposed that binding of MacP to PBP2a may result in a shift from an inactive to an active state, potentially through modulation of the conformation of the jaw subdomain^21^. It may therefore be proposed that the amino acid substitution A124T in PBP1a could enhance PBP1a activity by inducing similar changes in the conformation of the jaw subdomain. In contrast to PBP1a, variants of PBP2a that do not require MacP are predominantly located at the interface between the jaw domain and the TM segment of PBP2a^21^. This indicates that MacP and GarP may have different activation pathways that ultimately result in the induction of analogous conformational changes, thereby activating their respective aPBPs. Altogether, it seems that bacteria have evolved a range of molecular mechanisms, which are likely related to their specific PG sacculus growth mode, cell shape or cell envelope organization, in order to facilitate the necessary conformational changes for the activation of the TP and/or GT activities of aPBPs.

To uncover the function of GarP, we used the laboratory pneumococcal strain R6. This strain is derived from a virulent serotype 2 isolate D39 and is widely used to study fundamental cellular processes, such as transformation, cell division and PG biosynthesis^45^. D39 cells are significantly wider than R6 cells, and morphological changes between the two strains have been linked to mutations in the gene encoding PBP1a^45,46^. Moreover, additional mutations have been associated with several genetic interactions that differ between the R6 and its D39 progenitor. For example, the *mreCD* and *rodZ* genes are essential in the D39 strain, but not in the R6 strain^47,48^. MreC, MreD and RodZ have been associated with the peripheral PG synthesis machinery (elongasome), particularly with the essential bPBP (PBP2b) and RodA^47,48^. The observation that null mutations in *pbp1a* can suppress the requirement for *mreCD* and *rodZ* in strain D39 suggests that MreC, MreD and RodZ may be required for the growth of D39 strains when PBP1a is functional, but are dispensable when PBP1a is inactivated or absent. Consistent with this, swapping the *pbp1a*^D39^ and *pbp1a*^R6^ alleles in strain R6 restored the requirement for MreC and MreD^48^. Therefore, expression of the *pbp1a*^R6^ allele suppresses the requirement for MreCD in the R6 genetic background. The *pbp1a*^D39^ and *pbp1a*^R6^ alleles differ by two mutations, notably the A^R6^124T^D39^ (A in R6 and T in D39) substitution in the GT domain. Strikingly, and as shown in Extended Data Table 2 and 3, the same A124T substitution is generated upon deletion of *pbp2a* in the *garP*_L68A-F71A_ genetic background. The A124T substitution most likely enhances PBP1a^R6^ activity since it partially compensates for the absence of GarP. From these observations, it can be inferred that PBP1a^D39^ may be more active and less dependent on GarP in the D39 background. In addition, it suggests that the regulation of PBP1a activity by GarP may be at play in the non-essentiality of MreCD and/or RodZ in the R6 genetic background. Thus, MreC, MreD and RodZ may be indispensable when PBP1a is active, or PBP1a may become toxic in the absence of MreC, MreD or RodZ when its activity is no longer regulated by GarP. GarP may thus act as a fail-safe mechanism to prevent PBP1a from being constitutively active and detrimental to the function of the elongasome^33,47,48^.

In addition to reducing cell width, the deletion of *garP* also results in the formation of minicells (Fig. 2). In the absence of g*arP*, we observed that septa can form at abnormal positions between the parental and equatorial division sites, giving rise to daughter cells of varying sizes (Fig. 2). In some cases, septa appear in such close proximity to the parental division site that they do not result in minicell formation, or lead to minicells that are barely detectable. Interestingly, minicells have also been observed in the *pbp1a* mutant in numerous studies, although they were never explicitly identified as such, suggesting that they were considered as anecdotal observations^48^. Furthermore, the formation of minicells in the absence of *pbp1a* appears to increase in a context where PBP2a is constitutively active (*pbp2a* A77T)^21^. These observations collectively suggest that the positioning of the division site, and consequently the binding of MapZ to its specific PG motif^30^, may be influenced when the activity of aPBPs is dysregulated, and that PG remodeling by aPBPs is required for the correct positioning of the division site (cell equator) of the daughter cells. Finally, the deletion of *pbp1a* strongly induces the expression of the WalRK two-component system regulon, and the WalK histidine kinase is required for the growth of Δ*pbp1a* cells^33^. A number of genes encoding PG hydrolases, such as the septum splitting hydrolase *pcsB*^26^, are under the control of WalRK. Overall, aPBPs and in particular PBP1a, may exert a direct or indirect influence on PG synthesis, repair and/or remodeling, which in turn affects the placement and the splitting of the septum.

This work has also revealed that GarP contains the consensus motif SRxxR, which mediates its interaction with the cell division protein GpsB (Fig. 7). This motif was initially described in division proteins from *Bacillus subtilis*^17^. In particular, the SRxxR consensus motif of three proteins (PBP1, YrrS and YpbE) has been shown to mediate their interaction with GpsB. The function of YpbE and YrrS remains unclear. However, it has been demonstrated that YrrS interacts with the sporulation- specific bPBP PBP4b^17^. Consequently, it has been proposed that both proteins, which are bridged by GpsB, play a role in spore formation^17^. Interestingly, YpbE exhibits a C-terminal LysM domain and possesses a predicted unfolded region as observed in GarP (Extended Data Fig.7). YpbE might thus represent a functional homolog of GarP, capable of stimulating one of the three *B. subtilis* aPBPs PBP2c, PBP2d and PBP4, which lacks an SRxxR motif. In contrast to YpbE and YrrS, the function of PBP1 has been extensively investigated, showing its key contribution to both the elongation and division machineries. In a pioneering study, GpsB was shown to direct PBP1 shuttling between the divisome and the elongasome^49^. More recently, the characterization of the hexameric organization of GpsB in *B. subtilis* has led to propose that GpsB may interact simultaneously with multiple PBP1 molecules, thereby controlling their spatial arrangement in the membrane and consequently their PG polymerization and cross-linking activities^17^. Interestingly, akin to *B. subtilis* PBP1, the pneumococcal PBP2a also exhibits an SRxxR motif that mediates its interaction with GpsB^17^. It can be thus proposed that GpsB may be responsible for the spatial organization of PBP2a in the pneumococcal membrane. PBP2a is also regulated by a specific activator, MacP, which lacks an SRxxR motif^20,21^. We thus propose a model in which the molecular dialog between the two aPBP/activator pairs (PBP1a/GarP and PBP2a/MacP) and GpsB allows proper remodeling and/or repair of the PG layer during the cell cycle (Fig. 8). Moreover, we have demonstrated that the SRxxR motif is also present in the PG deacetylase PgdA and the muramidase MpgA. Interestingly, our findings also indicate that the sensitivity to lysozyme, which is known to be increased by the absence of PgdA^36^, is also increased by the absence or inactivation of PBP1a or GarP, and that *pgdA* plays a critical role in the synthetic relationship between *mpgA* and *pbp1a* or *garP* (Extended Data Table 4). Altogether, these observations suggest that the hydrolytic activity of PgdA may be linked to the activity of PBP1a, and that MpgA may require precise coordination with that of PgdA. Supporting this, PgdA activity is also regulated by GpsB and PBPA1 in *Listeria monocytogenes*^50^. In conclusion, the PBP1a/GarP and PBP2a/MacP pairs may work with MpgA, PgdA and GpsB to remodel and/or repair the PG layer (Fig. 8). The hexameric organization of GpsB provides six binding sites for the aforementioned proteins with the SRxxR motif. A challenging work is now required to elucidate the stoichiometry of these interactions, their assembly and disassembly dynamics, as well as possible binding competitions to determine whether they form a unique complex or several sub-complexes.

**Figure 8:**
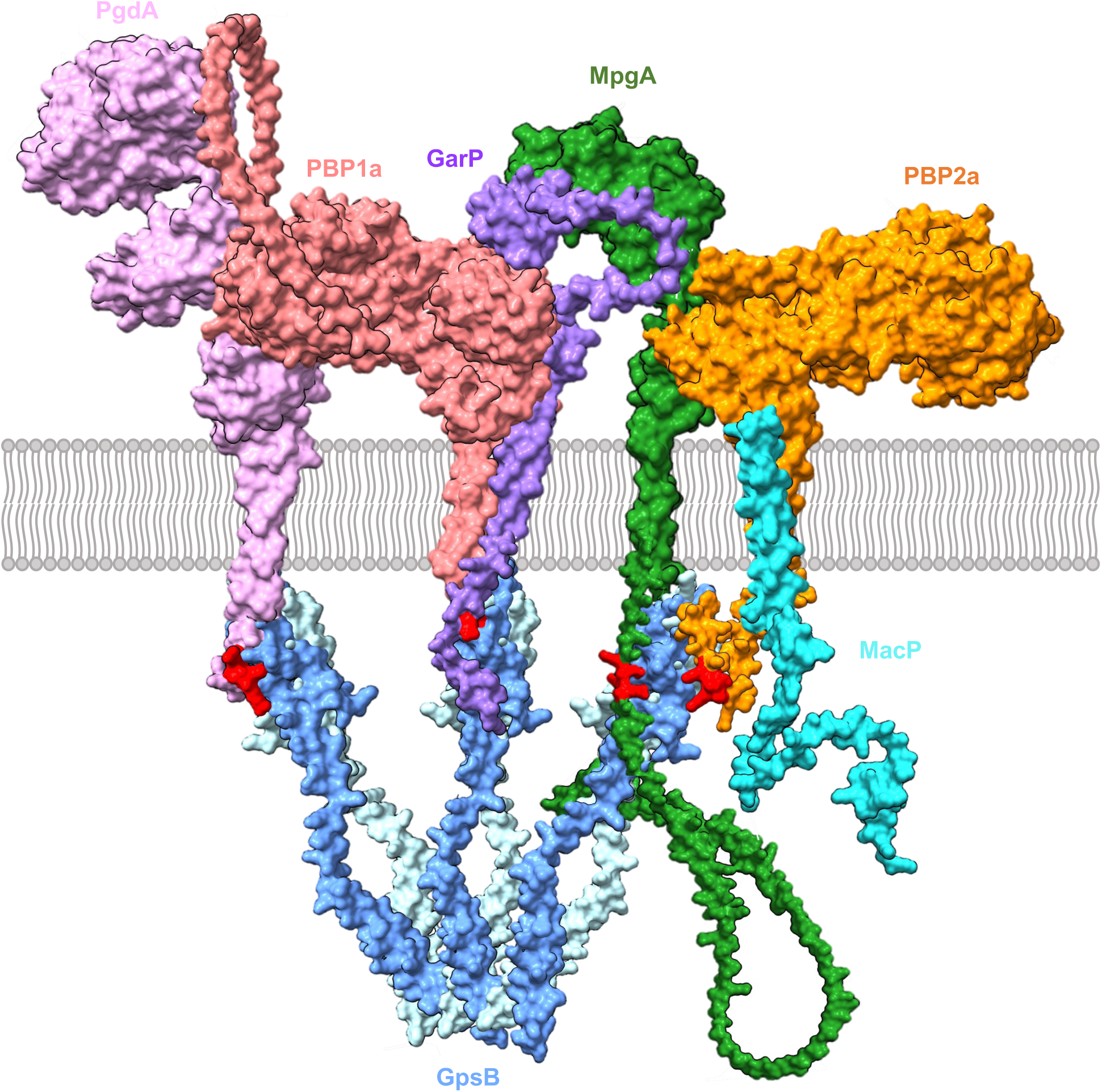
Model of the organization of the cell wall remodeling complex of *S. pneumoniae.* GarP (purple) modulates the activity of PBP1a (salmon) while MacP (cyan) regulates the activity of PBP2a (orange). PgdA (N-acetylglucosamine deacetylase), MpgA (muramidase) and GpsB are shown in pink, green and blue, respectively. The SRxxR motif of GarP, PBP2a, MpgA and PgdA a is represented in red. The interaction between SRxxR motifs and GpsB as well as those between GarP and PBP1a, and between MacP and PBP2a, are validated experimentally and are modeled using Alphafold3. The interaction surfaces between MpgA and PBP2a, MpgA and GarP and PgdA and PBP1a are not validated experimentally and not based on Alphafold modeling. The model does not presume any particular stoichiometry, dynamics, or binding competitions between the proteins involved.

In light of this model, a series of observations and hypotheses will be subjected to further investigation to understand the molecular pathways required for the biosynthesis and remodeling of the cell wall. This is particularly relevant in the context of the relationship with other regulators, such as the protein CopD and CozEa, which influence the spatiotemporal localization of PBP1a^18,19^. Moreover, several proteins within this complex, including GpsB, MpgA and MacP, have been shown to be phosphorylated by the membrane S/T-kinase StkP. More precisely, PBP2a activity is dependent upon MacP phosphorylation^20^, while GpsB controls the localization and activity of StkP^51^. GpsB has also been shown to be phosphorylated by the S/T-kinase PrkC and to regulate PrkC activity through a negative feedback loop in *B. subtilis*^52^. Importantly, GpsB is a paralog of DivIVA. The latter interacts with both GpsB and PBP1a, as well as PBP2a, and this interaction does not involve binding to the SRxxR motif^10,53^. While GpsB is required for pneumococcal cell constriction, as cells devoid of *gpsB* elongate, DivIVA is required for cell elongation^51^. DivIVA has also been recently shown to influence peripheral PG synthesis and the contribution of PBP1a and PBP2a to cell elongation during the early stage of the pneumococcal cell cycle^54^. DivIVA is also phosphorylated by StkP in *S. pneumoniae*.

Together with GpsB, it represents a rare example of division proteins phosphorylated in a variety of bacteria, including *B. subtilis*, *Mycobacterium tuberculosis and Streptomyces coelicolor*^10^. A deeper understanding of the molecular interplay between GpsB and DivIVA, along with the role of their phosphorylation state and that of MpgA and MacP, will be crucial to establish a comprehensive model and further elucidate the inner workings of this complex, particularly with regard to its stoichiometry and assembly dynamics, as well as is contribution to PG remodeling and repair in *S. pneumoniae*.

## METHODS

### Strain and growth conditions

*Streptococcus pneumoniae* R6 strains and its derivatives (Table S1) were cultured in C + Y or Todd Hewitt Yeast broth (THY) at 37°C. The optical density (OD)_550nm_ was acquired automatically in a Tecan SUNRISE microtiter plate every 10 min. Bacteria were grown on THY 1% agar plates supplemented with 3% (v/v) horse blood defibrinated. For antibiotic selections, THY plates were supplemented with kanamycin, streptomycin or chloramphenicol at final concentration 200 μg/mL, 250 μg/mL, and 2.5 μg/mL respectively. Strains expressing genes under the pComX promoter were cultured at 37°C in presence of the inducer peptide ComS^55^ at desired concentration. Pneumococcal mutant strains (gene deletions, ectopic gene expression, fluorescent protein fusions) were constructed by homologous recombination, using a two-step procedure, based on a bicistronic cassette called Janus^56^. All the strains were constructed by transformation of the WT strain or derivatives with a PCR amplicon, as previously described^57^. Primers and Templates used for the generation of amplicons are listed in (Table S1 and S2). The *E. coli* XL1-Blue, DH5α and TOP10 strains were used for cloning, *E. coli* BTH101 for bacterial two hybrid (BACTH) and *E. coli* BL21 (DE3) for protein overexpression. *E. coli* cells were grown in Luria Bertani broth (LB) supplemented with appropriated antibiotics.

### Transformation assays

Transformation was performed by homologous recombination as previously described^57^. All amplicons containing around 0.5-1 kb of flanking chromosomal DNA for homologous recombination were obtained by PCR using primers listed in Table S2. Recipient strains were grown until OD_550nm_ =0.1 and 3 μL of CSP-1 (competence stimulatory peptide, type 1) was added to 1ml of culture to induce competence. The mixture was incubated at 37°C for 10 min. 50 ng of purified amplicon was added to 50 μL transformation mixture and incubated at 37°C for 30 min. 100 μL of the mixture was spotted in plate with THY 1% agar supplemented with 3% (v/v) horse blood defibrinated and appropriated antibiotics (Kanamycine 0.25 mg/ml). Transformants were incubated overnight at 37°C for 20–24 h, at which time colony numbers were counted. All transformations were performed with *ΔcbpM::kan-rpsL* amplicons as a positive control and no DNA as negative control.

### Bacterial two hybrid

The BACTH (Bacterial Adenylate Cyclase Two-Hybrid) system kit was used according to the manufacturer’s protocol (Euromedex). Each pair of plasmids was co-transformed and co-transformants were re-streaked on LB agar plates supplemented with ampicillin 0.1 mg/ml, kanamycin 0.05 mg/mL, isospropyl-β-D-thiogalactopyranoside (IPTG) 0.5 mM, and X-gal 100 μg/mL. Plates were incubated at room temperature and photographed after 24 h and every 5 h for PBP1a/GarP interactions and after 48 h for GpsB interactions to monitor the appearance of blue colonies. Plasmids used in this study are listed in Table S1.

### Phase contrast and fluorescence microscopy

Pneumococcal cells were grown until OD_550nm_ = 0.1/0.2 and 1 μL of cultures was spotted onto 1% agarose C+Y pad on a microscopy slide and covered with a cover glass. Slides were observed using a Nikon TiE microscope fitted with an Orca-CMOS Flash4 V2 camera with a 100×/NA 1.45 objective. Strains expressing genes under pComX promoter were cultured in presence of the inducer peptide ComS at desired concentration at 37°C, 30 min before observation. Time-lapse microscopy was performed with a thermostatic chamber at 37°C and images were captured every 3 min for wide field microscopy and every 1 s for TIRF microscopy. Fluorescence images were acquired with a minimal exposure time to minimize bleaching and phototoxicity effects. Images were collected using NIS-Elements (Nikon) and analyzed using the software ImageJ (http://rsb.info.nih.gov/ij/) and the plugin MicrobeJ^58^ to generate fluorescent images, violin plots and kymographs. Statistical tests and details can be found in the figure legends.

### 3D-SIM and SIM Microscopy

Pneumococcal cells were grown until OD_550nm_ = 0.1/0.2 and visualized using a Zeiss Elyra 7 / Lattice SIM microscope equipped with a 63×/NA1.46 oil immersion objective and a pco.edge 4.2 sCMOS camera (Photon Lines), 405 nm, 488 nm, 561 nm, and 642 nm lasers for excitation.

For 3D-SIM imaging, each 3D-SIM stacks (512 x 512 pixels) consisting of 21 z-sections separated by 100-nm z-steps were imaged in SIM mode (13 phases). Raw images were SIM-processed and channel-aligned using Zen Black software (Zeiss) and observed with ImarisViewer software (https://imaris.oxinst.com/imaris-viewer).

### Scanning Electron Microscopy

Pneumococcal cells were grown until OD_550nm_ = 0.1/0.2 and then centrifugated at 1,000 g for 5 min. Pellets were resuspended in 1 ml of PBS and deposited on dried 12 mm diameter slides that were previously soaked in a poly-L-lysine solution (0.1 mg/ml) for 5 min with gentle agitation, then rinsed with water and dried for 2 h. After 10 min of incubation, the slides were washed three times (10 min) with cacodylate 0.18 M (pH 7.2) and incubated overnight with 2% glutaraldehyde in cacodylate 0.1 M (pH 7.2). The slides were then washed three times (10 min) with cacodylate 0.18 M (pH 7.2) and dehydrated by successive incubations in 30, 50, 70, 95 and 100% ethanol solution for 10 min. The slides were then incubated three times in Hexamethyldisilazane (HMDS) for 10 min and then dried on an absorbent paper. Samples were then mounted on holders with silver glue and metallized prior to observation. Samples were examined with a Zeiss Merlin Compact VP microscope fitted with an EBSD oxford Symmetry S3 camera (Oxford Intruments). Images were collected with Zeiss SmartSEM (Zeiss).

### Peptidoglycan labelling with fluorescent D-amino-acids

Pneumococcal cells were grown until OD_550nm_ = 0.1/0.2 and incubated for 5 min (short-pulse) or 3 h (long-pulse) at 37°C in C+Y with 500 mM of TAMRA-D-lysine. Cultures were washed three times with 1 ml of cold C+Y and then resuspended in 50 μL cold C+Y before observation.

### Co-immunoprecipitation

Pneumococcal cells were grown in THY at 37°C until OD_550nm_ = 0.4 and then centrifugated at 5,000 g for 15 min. Pellets were resuspended in a wash buffer (20 mM Tris-HCl pH 7.5, 200 mM NaCl) and centrifugated at 5,000 g for 10 min. Pellets were resuspended in a protoplast buffer (100 mM Tris-HCl pH 7.5, 2 mM MgCl_2,_ 1M Sucrose, 6 μg/mL of DNAse/ RNAse, 1x protein inhibitor Roche, 800 U mutanolysin, 8 mg/ml lysozyme) and incubated at 30°C for 30 min, and then centrifugated at 5,000 g for 10 min. Pellets were resuspended in a hypotonic buffer (100 mM Tris-HCl pH 7.5, 100 mM NaCl, 1 mM EDTA, 1 mM DTT, 1 mM MgCl_2_, 1% digitonin, 6 μg/mL of DNAse/RNAse, 1x protein inhibitor Roche, 8 U mutanolysin, 8 mg/mL lysozyme) and incubated at 37°C for 30 min. After centrifugation at 15,000 g for 30 min at 4°C, the supernatant was incubated with 35 μL of GFP-TRAP resin (Chromotek) at 4°C for 2 h under agitation. Protein-bound GFP-TRAP resin were eluted with dilution buffer (10 mM Tris-HCl pH 7.5, 150 mM NaCl, 0.5 mM EDTA, 1x protein inhibitor Roche, 0.05% digitonin) and analyzed by SDS-PAGE and immunoblot with either anti-GFP and anti-PBP1a antibodies.

### Preparation of *S. pneumoniae* crude extracts and immunoblot analysis

Pneumococcal cells were grown in C+Y medium until OD_550nm_ = 0.3-0.5, and then centrifugated at 5,000 g for 5 min. Pellets were resuspended in a lysis buffer (Tris-HCl 10 mM pH 8, EDTA 1 mM supplemented with 1:100 protease inhibitors cocktail). After cell disruption by sonication, non-lysed cells were discarded by centrifugation and crude extracts were normalized, analyzed by SDS-PAGE and transferred onto an immobilon-P membrane (Millipore). Primary antibodies were used at 1:5,000 (anti-GFP, AmsBio) in TBST-BSA 1%, 1:20,000 (anti-PBP1a^59^) in TBST-BSA 1%, 1:1,000 (anti-FLAG, Sigma) in PBS-milk 3% and 1:250,000 (anti-enolase^60^) in TBST-BSA 5%. The goat anti-rabbit or anti-mouse secondary antibody HRP conjugate was used at 1:5,000 (Biorad) in TBST-BSA 1%. The revelation was performed with Chemiluminescent SuperSignal West Pico PLUS kit (ThermoScientific) and imaged with Fusion Fx7 camera (Vilber Lourmat).

### Protein purification and preparation of glutamate- and iso-glutamine-containing lipid II

The recombinant extracellular region of PBP1a from strain R6 spanning residues 31 to 719 and the recombinant complex amidotransferase MurT/GatD were produced and purified as described previously^34,61^. MurT/GatD, the amidotransferase required for the amidation of glutamate into D-iso-glutamine prior PG cross-linking by pneumococcal PBPs Lys-containing lipid II, and dansylated lipid II were produced as described previously^62,63^. GarP full length was purified using the 6 histidine-tag encoded by the pt7-7 plasmid described above. Recombinant plasmid overproducing GarP full length was transformed into BL21 (DE3) *E. coli* strain. Cells were grown at 37°C until OD_600nm_ = 0.5 and gene expression was induced with 0.5 mM IPTG overnight at 25°C. Cells were harvested by centrifugation at 5,000 g for 10min at 4°C and resuspended in buffer A (50 mM HEPES pH 7.5, 150 mM NaCl, 1 mM MgCl_2_, DDM 1%, glycerol 10%) supplemented with 6 μg/mL DNAse/RNAse, 8 mg/mL lysozyme and 1:100 protease inhibitors cocktail. After sonication and centrifugation at 30,000 g for 30 min, soluble proteins were incubated with a Ni-NTA column (Qiagen) for 30 min, and washed with buffer A’ (50 mM HEPES pH 7.5, 150 mM NaCl, 1 mM MgCl_2_, DDM 0.1 %, glycerol 10 %) and buffer B (50 mM HEPES pH 7.5, 150 mM NaCl, 1 mM MgCl_2_, 20 mM imidazole, DDM 0.1%, glycerol 10 %). Elution was performed with elution buffer (50 mM HEPES pH 7.5, 150 mM NaCl, 1 mM MgCl_2_, 300 mM imidazole, DDM 0.1 %, glycerol 10 %). Elution fractions were analyzed by SDS-PAGE and the fractions containing proteins were pooled and dialyzed with dialysis buffer (50 mM HEPES pH 7.5, 150 mM NaCl, 1 mM MgCl_2_, DDM 0.1 %, glycerol 10 %). The dialyzed sample was then gel filtrated using a Superdex200 10/300 increase column (Cytiva) equilibrated with dialysis buffer. Pure proteins were concentrated and stored at -80°C.

### *In vitro* reaction of peptidoglycan synthesis

The required amount of lipid II mixture was dried and redissolved in the appropriate volume of 12 µM MurT/GatD, 20 mM L-glutamine, 20 mM ATP in 50 mM HEPES, pH 7.5, 150 mM NaCl, 1 mM MgCl_2_. After overnight incubation at 30°C, DMSO, PBP1a with or without GarP were added to have the following final concentrations and conditions: 50 µM unlabeled lipid II, 5 µM dansylated lipid II, 1 µM PBP1a, 6 µM GarP, in 45 mM HEPES, pH 7.5, 135 mM NaCl, 0.9 mM MgCl_2_, 0.009% triton X-100, 25% DMSO. The reaction mix also contained 5 mM ATP, 5 mM L-Gln and 3 µM MurT/GatD carried over from the lipid II amidation reaction. After incubation at 37°C for various time intervals, aliquots were withdrawn and the reactions were stopped by addition to an equal volume of 10 mM penicillin G and 10 mM moenomycin in water. Samples were analyzed by polyacrylamide gel electrophoresis^64^. Gels were imaged under UV-transillumination with a Bio-Rad ChemiDoc XRS+ instrument. The in gel fluorescent intensities were quantified from non-saturated tif images using the ImageJ software.

### Structural Predictions

Structural predictions of GarP from protein sequence were performed using Alphafold3 (https://alphafoldserver.com), and MobiDB (https://mobidb.org). Structural prediction of GarP, PBP1a, GarP/PBP1a, YpbE and remodeling complex were run using alphafold3. Model visualization and figure preparation were performed using ChimeraX software (https://www.cgl.ucsf.edu/chimerax/).

## AUTHOR CONTRIBUTION

HM, AD, and CG designed research; HM, CL, AD, CF, CC and AZ performed experiments; HM, AD, AZ, CM and CG analyzed data; HM, AD, and CG wrote the manuscript; AD and CG jointly supervised the work. All authors revised the manuscript.

### ACKNOWLEDGMENTS

We thank Nathalie Campo and Eefjan Breukink for providing us with the silica micropillar mold for the preparation of microhole agarose pads and Lipid II, respectively. Support for this work comes from the CNRS, the Université Lyon I, the foundation Bettencourt-Schuller to CG and the Agence Nationale de la Recherche (ANR-23-CE11-0029 and ANR-19-CE15-0011 to CM and CG and ANR-20-CE07- 0012 to AZ). We acknowledge the contribution of the microscopy facility (PLATIM) of the SFR Biosciences (Université Claude Bernard Lyon 1, CNRS UAR3444, INSERM US8, ENS). IBS acknowledges integration into the Interdisciplinary Research Institute of Grenoble (IRIG, CEA).

## EXTENDED DATA FIGURES

**Extended Data Fig. 1:**
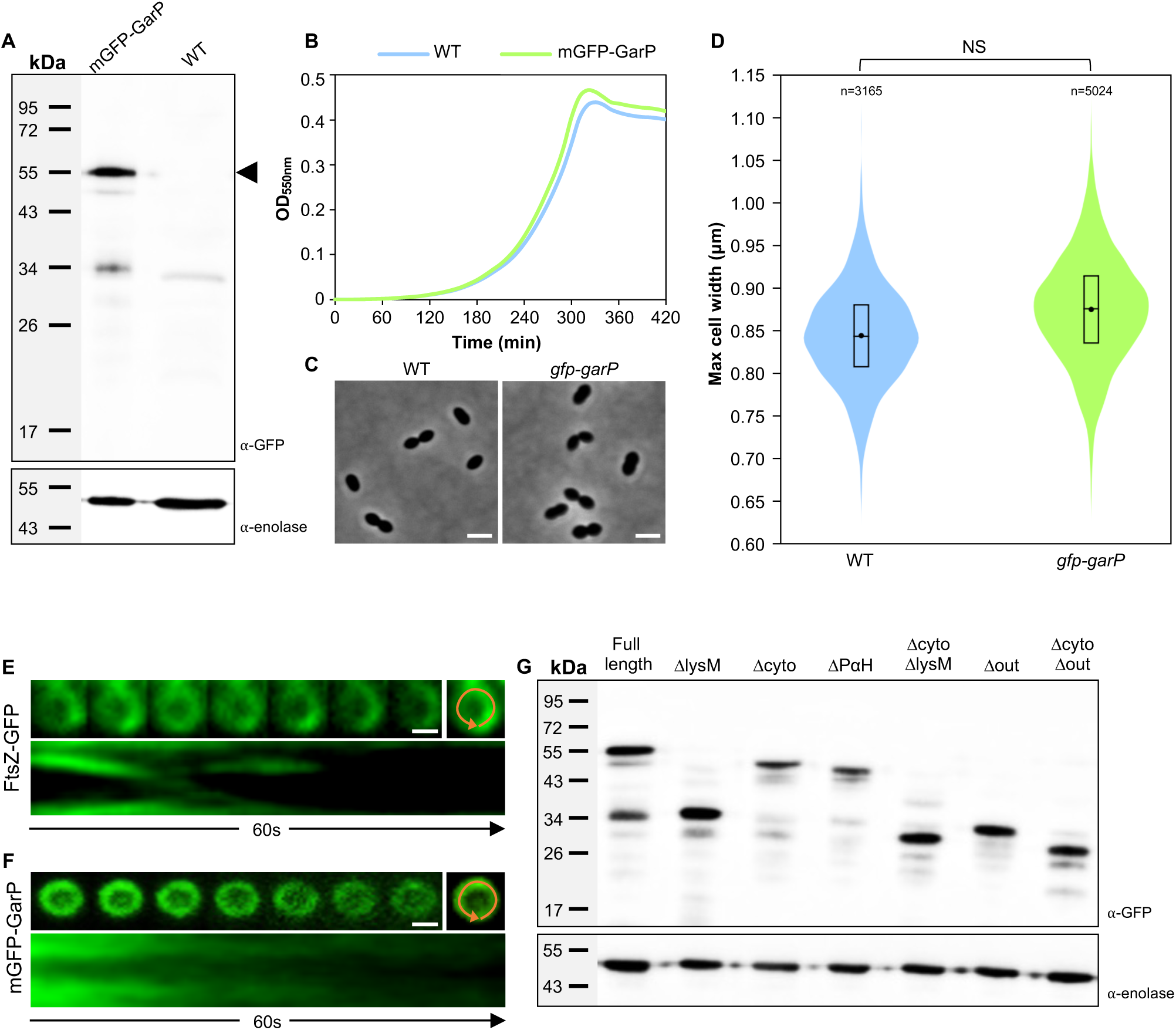
GarP is a membrane protein that forms a ring at midcell. **(A)** Western immunoblot of whole-cell lysates of wild-type (WT) and *gfp-garP* strains with specific anti-GFP antibody and anti-enolase antibody as a loading control. The expected mGFP-GarP band is indicated by the black triangle. **(B)** Growth of WT and *gfp-garP* strains in C+Y media at 37°C. **(C)** Phase contrast microscopy images of WT and *gfp-garP* strains. Scale bars, 2 µm. **(D)** Violin plots showing the distribution of the maximum cell width from three independent experiments, with n indicating the total number of cells. The box indicates the 25^th^ to the 75^th^ percentile. The mean and the median values are indicated with a dot and a line in the box, respectively. Statistical comparison was done using a t-test. NS, not significant (p > 0.05). **(E-F)** Representative montage of images showing the dynamics of FtsZ-GFP (E) or mGFP-GarP (F) in WT cells observed by conventional fluorescence microscopy in a microhole for 60 s (9 s intervals). A summation of all the images and a kymograph are shown on the right and below the montage respectively. The kymographs (1 frame/s) were generated from a circular line around the circumference of the cell, highlighted in orange on the summation images. Scale bars, 0.5 µm. **(G)** Western immunoblot of whole-cell lysates of mGFP-GarP fusion (Full length) and derivatives with specific anti-GFP antibody and anti-enolase antibody as a loading control.

**Extended Data Fig. 2:**
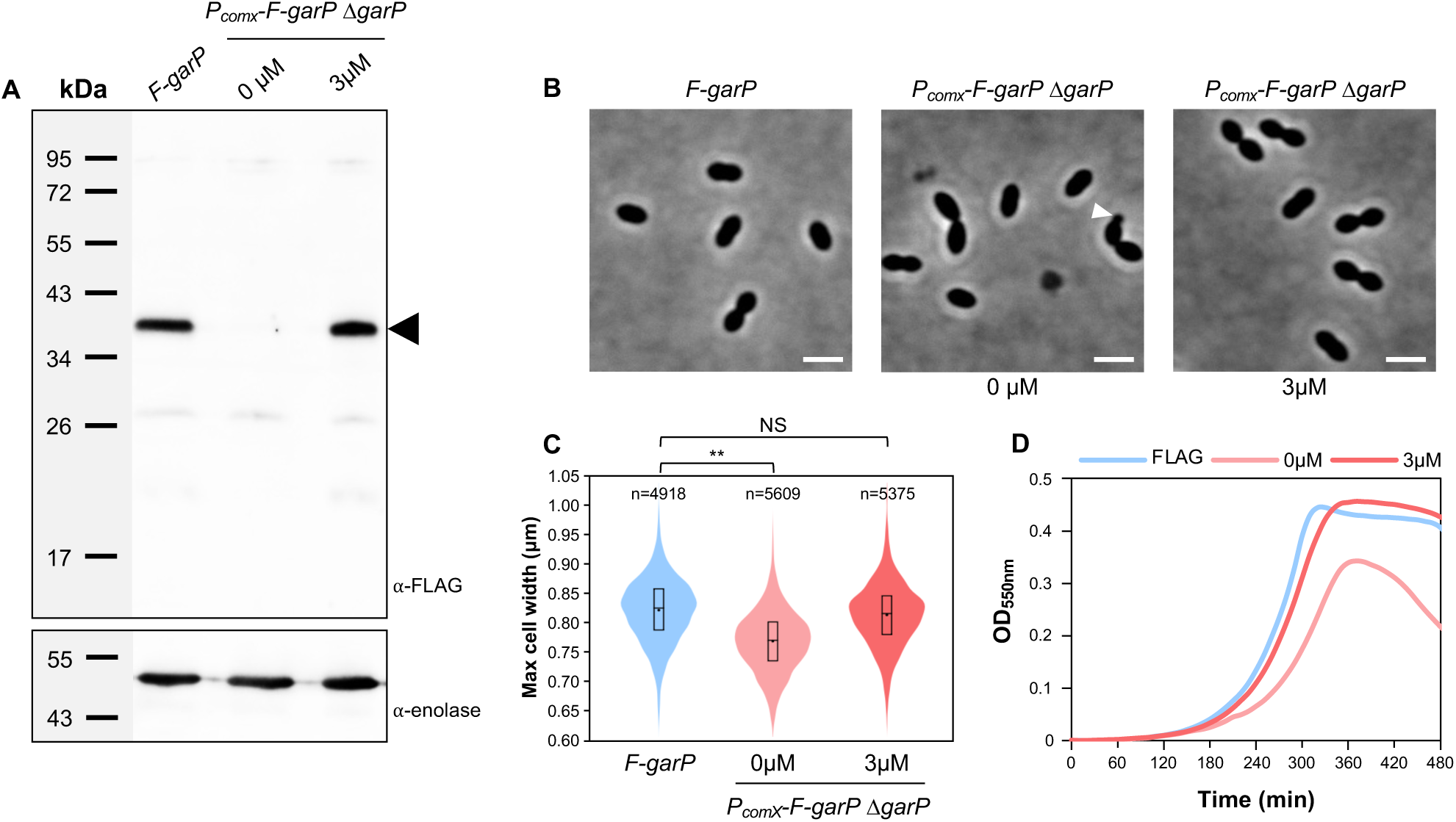
Expression, cell growth and cell morphology of *FLAG-garP* and *PcomX- FLAG-garP ΔgarP* cells. **(A)** Western immunoblot of whole-cell lysates of *FLAG-garP* and *PcomX-FLAG-garP ΔgarP* cells, grown in the presence (3 µM) or the absence (0 µM) of inducer (ComS), revealed with anti-FLAG antibody and anti-enolase antibody as a loading control. The expected band for FLAG-GarP is indicated by the black triangle. **(B)** Phase contrast microscopy images of *FLAG-garP* and *PcomX- FLAG-garP ΔgarP* cells, grown in the presence (3 µM) or absence (0 µM) of ComS inducer. Minicells are highlighted by a white triangle. Scale bars, 2 µm. **(C)** Violin plots showing the distribution of the maximum cell width from three independent experiments, with n indicating the total number of cells. The box indicates the 25^th^ to the 75^th^ percentile. The mean and the median are indicated with a dot and a line in the box, respectively. Statistical comparison was done using a t-test. NS, not significant, p >0.05. **(D)** Growth of *FLAG-garP* and *PcomX-FLAG-garP ΔgarP* in C+Y medium at 37°C.

**Extended Data Fig. 3:**
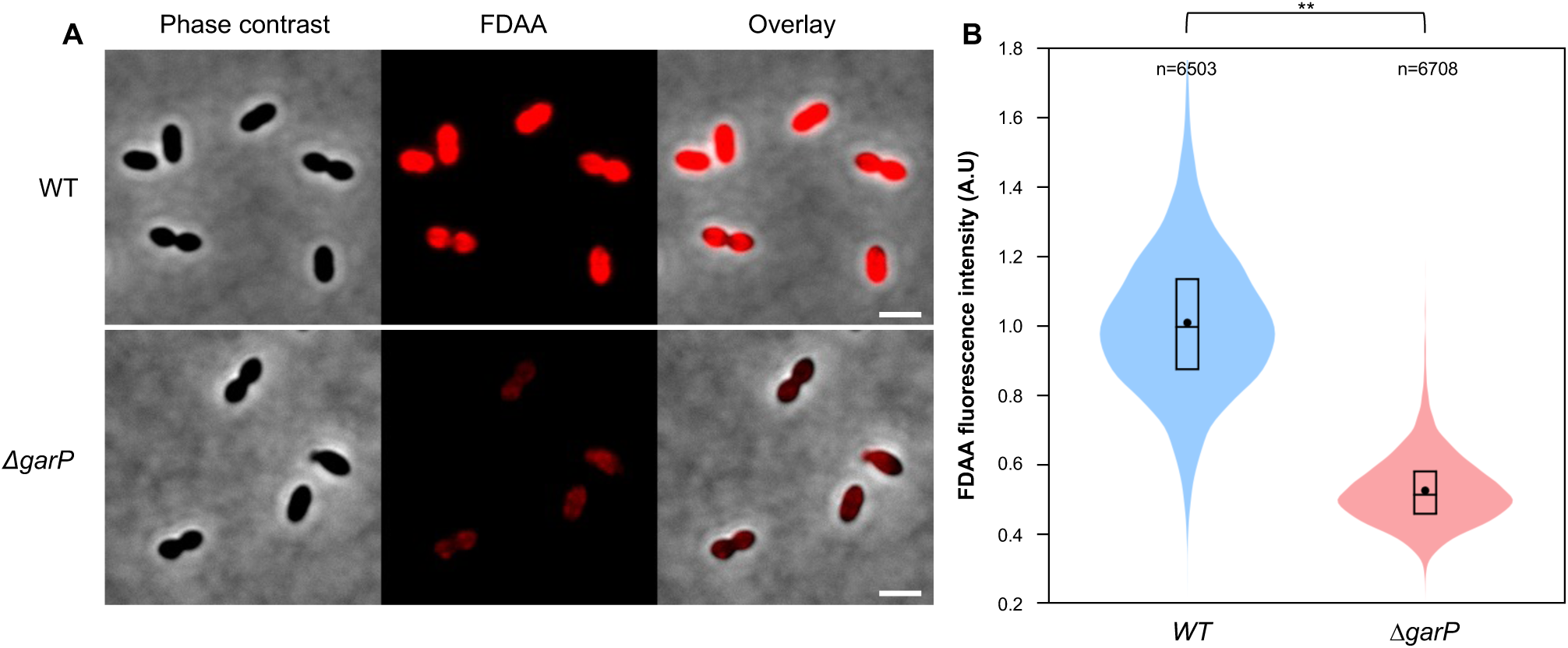
Minicell formation results from aberrant ectopic division septa and the absence of GarP leads to reduced FDAA incorporation. **(A)** Representative images of wild-type and *ΔgarP* cells after a long period of FDAA (TDL) labelling. The phase contrast, the RFP and the merged image between the RFP and the phase contrast channels are shown. Scale bars, 2 µm. **(B)** Violin plots showing the distribution of the normalized FDAA fluorescence intensity for wild-type and *ΔgarP* cells after a long period of FDAA (TDL) labelling from three independent experiments, with n indicating the total number of cells. The box indicates the 25^th^ to the 75^th^ percentile, and the whiskers indicate the minimum and the maximum values. The mean and the median are indicated with a dot and a line in the box, respectively. Statistical comparison was done using a t-test. **p < 0.01.

**Extended Data Fig. 4:**
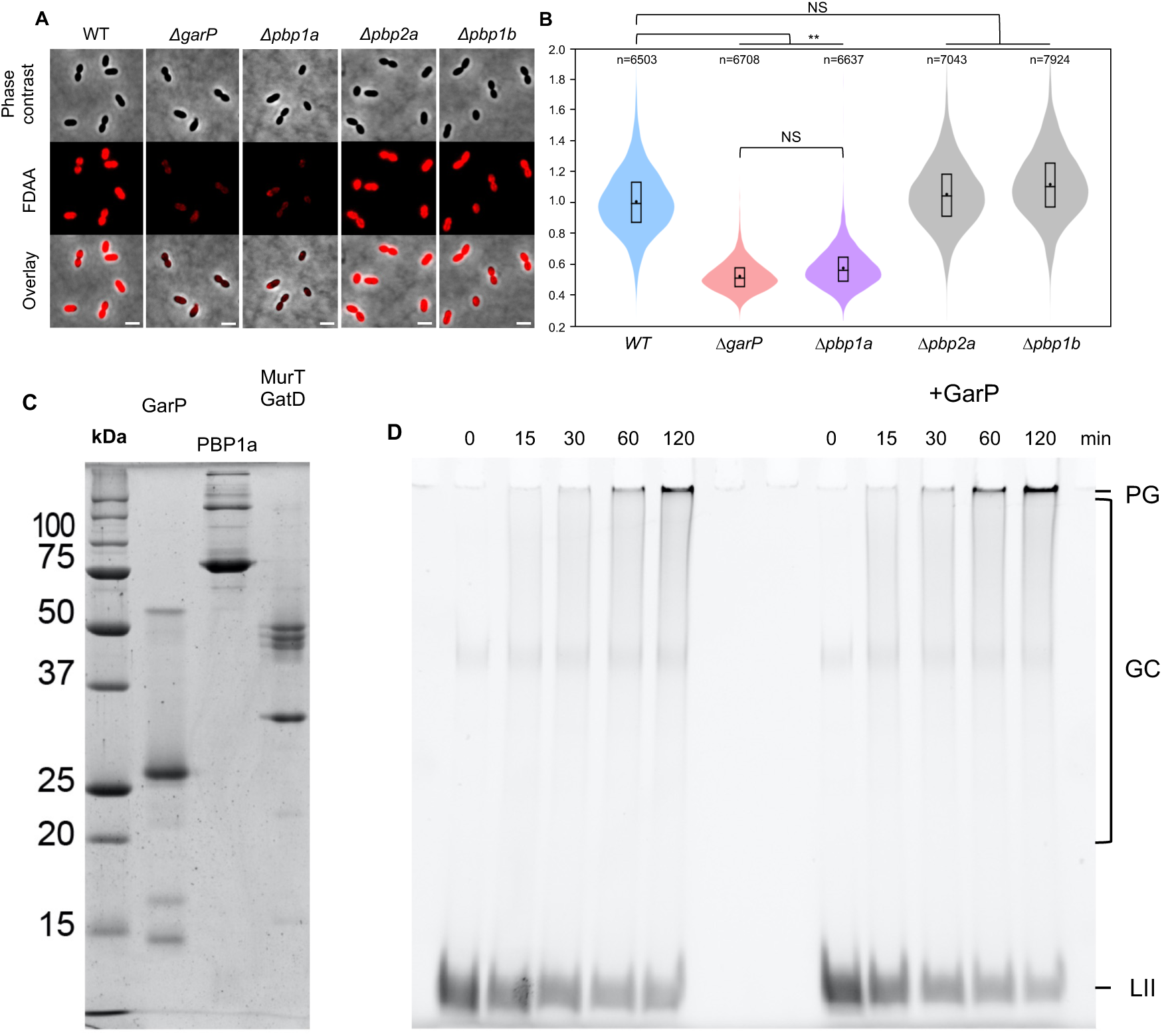
GarP interacts with and promotes the activity of PBP1a. **(A)** Representative images of wild-type (WT), *ΔgarP, Δpbp1a*, *Δpbp2a* and *Δpbp1b* cells after a long period of FDAA (TDL) labelling. The phase contrast, the RFP and the merged images between the RFP and the phase contrast channels are shown. Scale bars, 2 µm. **(B)** Violin plots showing the distribution of the normalized FDAA fluorescence intensity for wild-type, *ΔgarP, Δpbp1a*, *Δpbp2a* and *Δpbp1b* cells after a long period of FDAA labelling, from three independent experiments, with n indicating the total number of cells. The box indicates the 25^th^ to the 75^th^ percentile. The mean and the median are indicated with a dot and a line in the box, respectively. Statistical comparison was done using a t-test. **p < 0.01 and NS, not significant, p > 0.05. **(C)** Coomassie-stained SDS-PAGE of the recombinant proteins used this study (GarP, PBP1a, MurT/GatD). **(D)** Gel electrophoresis analysis of a time course of peptidoglycan assembly from a mixture of unlabeled and fluorescently labeled lipid II by PBP1a in the absence (left) and presence (right) of GarP. After various time intervals, the reaction was stopped by addition of penicillin G and moenomycin. The gel was imaged under UV- transillumination. Lipid II (LII) migrates at the front, uncross-linked glycan chains (GC) migrate as a smear within the gel, cross-linked peptidoglycan (PG) remains in the wells.

**Extended Data Fig. 5:**
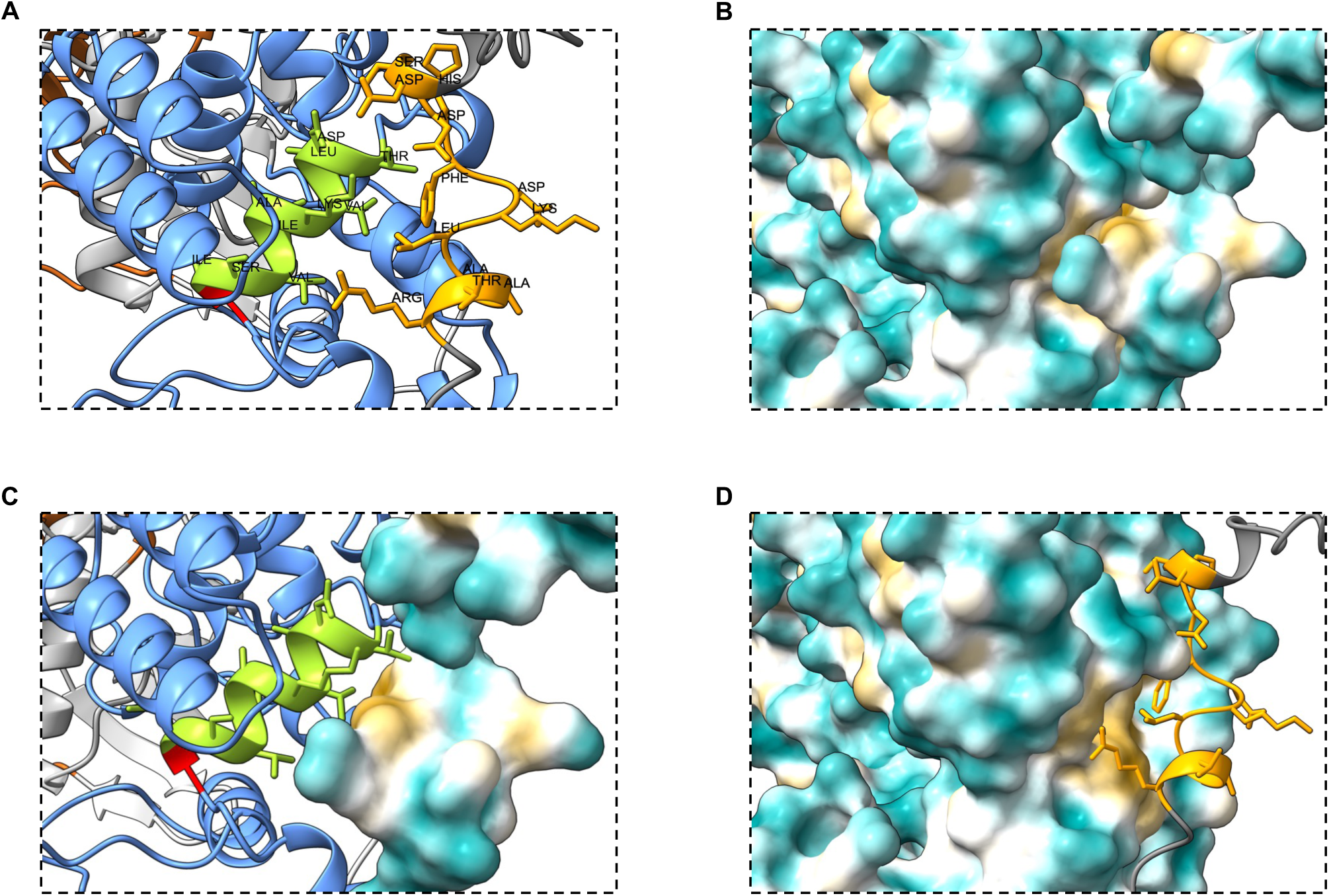
The PαH of GarP is required for PBP1a activation. (A-B) Different representations of the interface between the PαH of GarP and the helix of GT domain of PBP1a (light green) bearing the catalytic residue (E91) highlighted in red. (**A**) Transparent surface and ribbon representations for both molecules. The name of amino acids is indicated. **(B)** Hydrophobicity surface representation. **(C)** Hydrophobicity surface and ribbon diagram representation for GarP and PBP1a, respectively. **(D)** Ribbon diagram and hydrophobicity surface representations for GarP and PBP1a, respectively.

**Extended Data Fig. 6:**
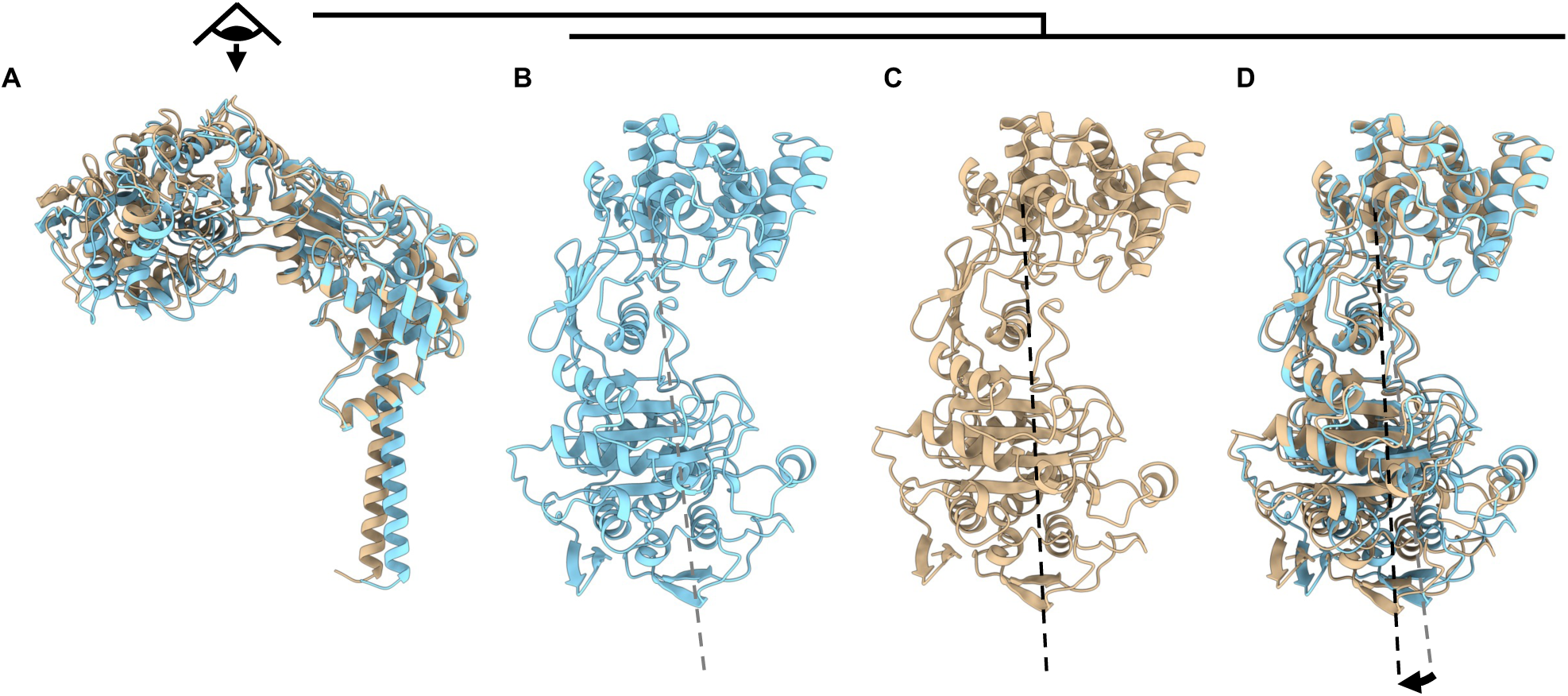
The PBP1a-A124T mutation bypasses the requirement for GarP. **(A)** Superposition between PBP1a alone (in light blue) and PBP1a in complex with GarP (in golden bronze), predicted by Alphafold3. Superposition is based on the GT domain. **(B)** Top view of PBP1a alone predicted by Alphafold3. The angle between the interdomain region and the GT domain is represented by the dotted line in grey. **(C)** Top view of PBP1a in complex with GarP predicted by Alphafold3. The angle between the interdomain region and the GT domain is represented by the dotted line in black. **(D)** Top view of the superposition between Alphafold3 models of PBP1a alone (in light blue) and PBP1a in complex with GarP (in golden bronze). The angle between the interdomain region and the GT domain for PBP1a alone or in complex with GarP is represented by the dotted line in grey and black, respectively. The deviation is represented by an arrow.

**Extended Data Fig. 7:**
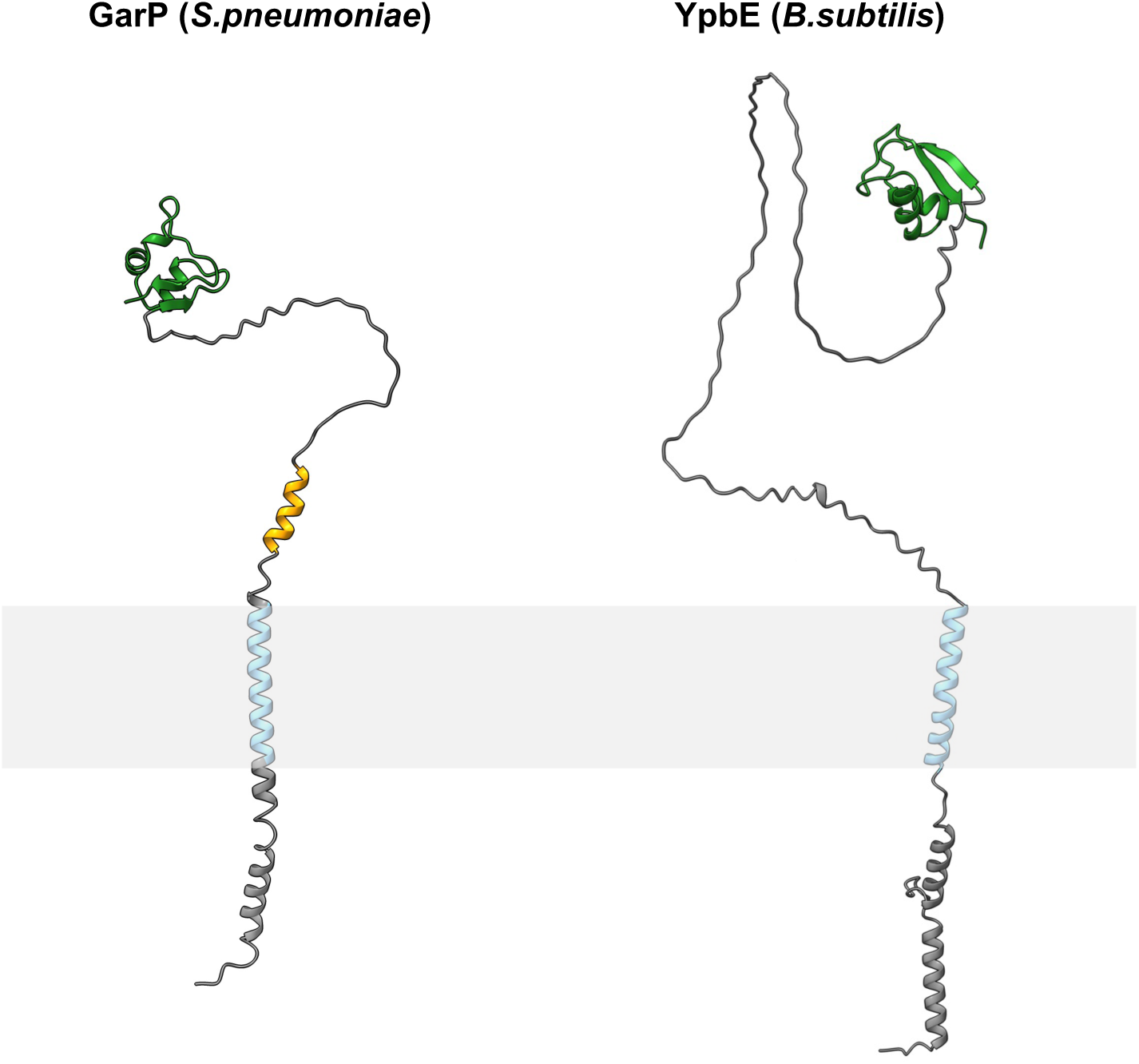
Structural homology between GarP (*S. pneumoniae*) and YpbE (*B. subtilis*). Alphafold3 models of GarP and YpbE. The predicted transmembrane segment and the LysM domain of both proteins are shown in blue and green, respectively. Intrinsically disordered regions are shown in gray. The predicted α-helix of GarP is shown in orange.

## EXTENDED DATA TABLES

**Extended Table 1.**
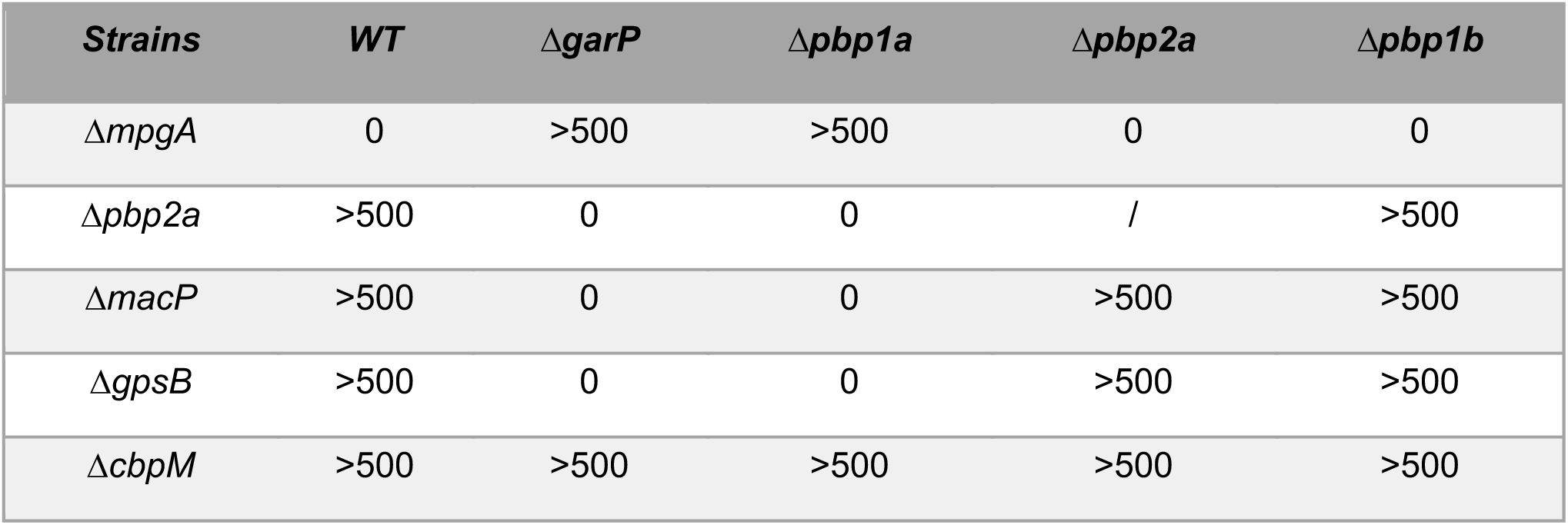
GarP interacts with and promotes the activity of PBP1a. Synthetic lethality and viability relationships. Results are presented as the number of colonies. Strains are indicated in white and amplicons in black. All transformation experiments were performed with no added DNA as the negative control and with a *ΔcbpM::kan-rpsL* amplicon as the positive control. Experiments made in triplicate.

1 EXTENDED DATA TABLES
| *Strains* | *WT* | *ΔgarP* | *Δpbp1a* | *Δpbp2a* | *Δpbp1b* |
| --- | --- | --- | --- | --- | --- |
| *ΔmpgA* | 0 | >500 | >500 | 0 | 0 |
| *Δpbp2a* | >500 | 0 | 0 | / | >500 |
| *ΔmacP* | >500 | 0 | 0 | >500 | >500 |
| *ΔgpsB* | >500 | 0 | 0 | >500 | >500 |
| *ΔcbpM* | >500 | >500 | >500 | >500 | >500 |

**Extended Table 2.**
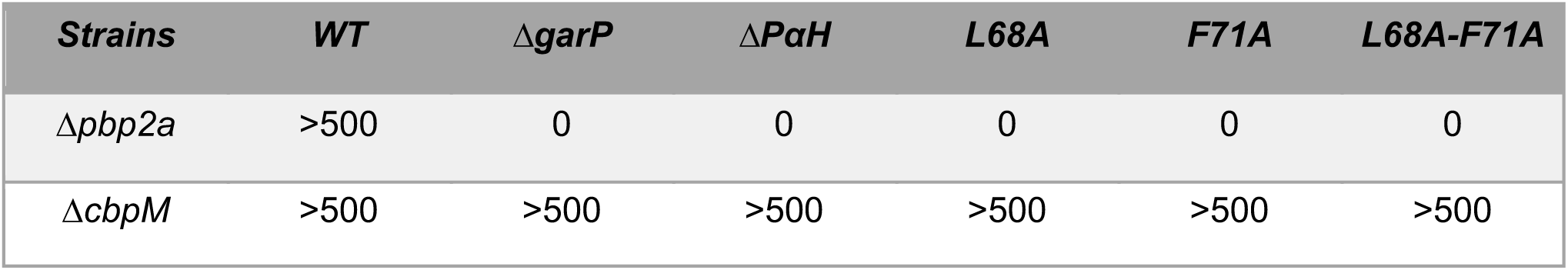
The PαH of GarP is required for PBP1a activation. Synthetic lethality and viability relationship. Results are presented as the number of colonies. Strains are indicated in white and amplicons in black. All transformation experiments were performed with no added DNA as the negative control and with a *ΔcbpM::kan-rpsL* amplicon as the positive control. Experiments made in triplicate.

**Extended Table 3.** *pbp1a-A124T* mutation compensates the lack of *garP*. Synthetic lethality and viability relationship. Results are presented as the number of colonies. Strains are indicated in white and amplicons in black. All transformation experiments were performed with no added DNA as the negative control and with a *ΔcbpM::kan-rpsL* amplicon as the positive control. Experiments made in triplicate.

| Strains | WT | $\Delta garP$ | *PBP1a-A124T* | *PBP1a-A124T*<br>$\Delta garP$ | *PBP1a-A124T*<br>L68A-F71A |
| --- | --- | --- | --- | --- | --- |
| $\Delta pbp2a$ | >500 | 0 | >500 | >500 | >500 |
| $\Delta cbpM$ | >500 | >500 | >500 | >500 | >500 |

**Extended Table 4.** A GpsB-binding motif SRxxR is conserved in GarP, PBP2a, PgdA and MpgA. Synthetic lethality and viability relationship. Results are presented as the number of colonies. Strains are indicated in white and amplicons in black. All transformation experiments were performed with no added DNA as the negative control and with a *ΔcbpM::kan-rpsL* amplicon as the positive control. Experiments made in triplicate.

| Strains | WT | $\Delta garP$ | $\Delta pbp1a$ | $\Delta pbp2a$ | $\Delta pgdA$ |
| --- | --- | --- | --- | --- | --- |
| $\Delta mpgA$ | 0 | >500 | >500 | 0 | >500 |
| $\Delta cbpM$ | >500 | >500 | >500 | >500 | >500 |

## REFERENCES

1. Egan, A. J. F., Errington, J. & Vollmer, W. Regulation of peptidoglycan synthesis and remodelling. Nat Rev Microbiol 18, 446–460 (2020).

2. Kumar, S., Mollo, A., Kahne, D. & Ruiz, N. The Bacterial Cell Wall: From Lipid II Flipping to Polymerization. Chem. Rev. 122, 8884–8910 (2022).

3. Sjodt, M. et al. Structural coordination of polymerization and crosslinking by a SEDS–bPBP peptidoglycan synthase complex. Nat Microbiol 5, 813–820 (2020).

4. Straume, D., Piechowiak, K. W., Kjos, M. & Håvarstein, L. S. Class A PBPs: It is time to rethink traditional paradigms. Molecular Microbiology 116, 41–52 (2021).

5. Massidda, O., Nováková, L. & Vollmer, W. From models to pathogens: how much have we learned about Streptococcus pneumoniae cell division? Environ Microbiol 15, 3133–3157 (2013).

6. Haeusser, D. P. & Margolin, W. Splitsville: structural and functional insights into the dynamic bacterial Z ring. Nat Rev Microbiol 14, 305–319 (2016).

7. Cho, H. et al. Bacterial cell wall biogenesis is mediated by SEDS and PBP polymerase families functioning semi-autonomously. Nat Microbiol 1, 1–8 (2016).

8. Perez, A. J. et al. Movement dynamics of divisome proteins and PBP2x:FtsW in cells of Streptococcus pneumoniae. Proc Natl Acad Sci U S A 116, 3211–3220 (2019).

9. Daniel, R. A. & Errington, J. Control of Cell Morphogenesis in Bacteria: Two Distinct Ways to Make a Rod-Shaped Cell. Cell 113, 767–776 (2003).

10. Hammond, L. R., White, M. L. & Eswara, P. J. ¡vIVA la DivIVA! Journal of Bacteriology 201, 10.1128/jb.00245-19 (2019).

11. Tsui, H.-C. T. et al. Pbp2x localizes separately from Pbp2b and other peptidoglycan synthesis proteins during later stages of cell division of treptococcus pneumoniae D39. Molecular Microbiology 94, 21–40 (2014).

12. Vigouroux, A. et al. Class-A penicillin binding proteins do not contribute to cell shape but repair cell-wall defects. eLife 9, e51998 (2020).

13. Typas, A. et al. Regulation of peptidoglycan synthesis by outer membrane proteins. Cell 143, 1097– 1109 (2010).

14. Egan, A. J. F. et al. Induced conformational changes activate the peptidoglycan synthase PBP1B. Molecular Microbiology 110, 335–356 (2018).

15. Sardis, M. F., Bohrhunter, J. L., Greene, N. G. & Bernhardt, T. G. The LpoA activator is required to stimulate the peptidoglycan polymerase activity of its cognate cell wall synthase PBP1a. Proceedings of the National Academy of Sciences 118, e2108894118 (2021).

16. Costa, S. F. et al. The role of GpsB in Staphylococcus aureus cell morphogenesis. mBio 15, e03235–23 (2024).

17. Cleverley, R. M. et al. The cell cycle regulator GpsB functions as cytosolic adaptor for multiple cell wall enzymes. Nat Commun 10, 261 (2019).

18. Fenton, A. K., El Mortaji, L., Lau, D. T. C., Rudner, D. Z. & Bernhardt, T. G. CozE is a member of the MreCD complex that directs cell elongation in Streptococcus pneumoniae. Nat Microbiol 2, 16237 (2016).

19. Lenoir, C. et al. The morphogenic protein CopD controls the spatio-temporal dynamics of PBP1a and PBP2b in Streptococcus pneumoniae. mBio 14, e01411–23 (2023).

20. Fenton, A. K. et al. Phosphorylation-dependent activation of the cell wall synthase PBP2a in Streptococcus pneumoniae by MacP. Proceedings of the National Academy of Sciences of the United States of America 115, 2812–2817 (2018).

21. Midonet, C. et al. MacP bypass variants of Streptococcus pneumoniae PBP2a suggest a conserved mechanism for the activation of bifunctional cell wall synthases. mBio 14, e02390–23 (2023).

22. Paik, J., Kern, I., Lurz, R. & Hakenbeck, R. Mutational Analysis of the Streptococcus pneumoniae Bimodular Class A Penicillin-Binding Proteins. Journal of Bacteriology 181, 3852–3856 (1999).

23. Rued, B. E. et al. Suppression and synthetic-lethal genetic relationships of ΔgpsB mutations indicate that GpsB mediates protein phosphorylation and penicillin-binding protein interactions in Streptococcus pneumoniae D39. Molecular Microbiology 103, 931–957 (2017).

24. Buist, G., Steen, A., Kok, J. & Kuipers, O. P. LysM, a widely distributed protein motif for binding to (peptido)glycans. Molecular Microbiology 68, 838–847 (2008).

25. Taguchi, A., Page, J. E., Tsui, H.-C. T., Winkler, M. E. & Walker, S. Biochemical reconstitution defines new functions for membrane-bound glycosidases in assembly of the bacterial cell wall. PNAS 118, (2021).

26. Barendt, S. M., Sham, L.-T. & Winkler, M. E. Characterization of Mutants Deficient in the l,d- Carboxypeptidase (DacB) and WalRK (VicRK) Regulon, Involved in Peptidoglycan Maturation of Streptococcus pneumoniae Serotype 2 Strain D39▿. J Bacteriol 193, 2290–2300 (2011).

27. Alcorlo, M., Martínez-Caballero, S., Molina, R. & Hermoso, J. A. Carbohydrate recognition and lysis by bacterial peptidoglycan hydrolases. Curr Opin Struct Biol 44, 87–100 (2017).

28. Fleurie, A. et al. MapZ marks the division sites and positions FtsZ rings in Streptococcus pneumoniae. Nature 516, 259–262 (2014).

29. Holečková, N. et al. LocZ Is a New Cell Division Protein Involved in Proper Septum Placement in Streptococcus pneumoniae. mBio 6, 10.1128/mbio.01700-14 (2014).

30. Manuse, S. et al. Structure-function analysis of the extracellular domain of the pneumococcal cell division site positioning protein MapZ. Nat Commun 7, 12071 (2016).

31. Kuru, E. et al. Mechanisms of Incorporation for D-Amino Acid Probes That Target Peptidoglycan Biosynthesis. ACS Chem. Biol. 14, 2745–2756 (2019).

32. Berg, K. H., Stamsås, G. A., Straume, D. & Håvarstein, L. S. Effects of low PBP2b levels on cell morphology and peptidoglycan composition in Streptococcus pneumoniae R6. J Bacteriol 195, 4342–4354 (2013).

33. Tsui, H.-C. T. et al. Suppression of a deletion mutation in the gene encoding essential PBP2b reveals a new lytic transglycosylase involved in peripheral peptidoglycan synthesis in Streptococcus pneumoniae D39. Mol Microbiol 100, 1039–1065 (2016).

34. Zapun, A. et al. In vitro Reconstitution of Peptidoglycan Assembly from the Gram-Positive Pathogen Streptococcus pneumoniae. ACS Chem. Biol. 8, 2688–2696 (2013).

35. Sung, M.-T. et al. Crystal structure of the membrane-bound bifunctional transglycosylase PBP1b from Escherichia coli. Proceedings of the National Academy of Sciences 106, 8824–8829 (2009).

36. Vollmer, W. & Tomasz, A. The pgdA Gene Encodes for a PeptidoglycanN-Acetylglucosamine Deacetylase in Streptococcus pneumoniae *. Journal of Biological Chemistry 275, 20496–20501 (2000).

37. Vollmer, W. & Tomasz, A. Peptidoglycan N-Acetylglucosamine Deacetylase, a Putative Virulence Factor in Streptococcus pneumoniae. Infection and Immunity 70, 7176–7178 (2002).

38. Straume, D. et al. Class A PBPs have a distinct and unique role in the construction of the pneumococcal cell wall. Proc Natl Acad Sci U S A 117, 6129–6138 (2020).

39. Höltje, J. V. Growth of the stress-bearing and shape-maintaining murein sacculus of Escherichia coli. Microbiol Mol Biol Rev 62, 181–203 (1998).

40. Egan, A. J. F. et al. Outer-membrane lipoprotein LpoB spans the periplasm to stimulate the peptidoglycan synthase PBP1B. Proceedings of the National Academy of Sciences of the United States of America 111, 8197–8202 (2014).

41. Lupoli, T. J. et al. Lipoprotein Activators Stimulate Escherichia coli Penicillin-Binding Proteins by Different Mechanisms. J. Am. Chem. Soc. 136, 52–55 (2014).

42. Caveney, N. A. et al. Structure of the Peptidoglycan Synthase Activator LpoP in Pseudomonas aeruginosa. Structure 28, 643–650.e5 (2020).

43. Sauvage, E. & Terrak, M. Glycosyltransferases and Transpeptidases/Penicillin-Binding Proteins: Valuable Targets for New Antibacterials. Antibiotics 5, 12 (2016).

44. Di Guilmi, A. M. et al. Identification, purification, and characterization of transpeptidase and glycosyltransferase domains of Streptococcus pneumoniae penicillin-binding protein 1a. J Bacteriol 180, 5652–5659 (1998).

45. Lanie, J. A. et al. Genome Sequence of Avery’s Virulent Serotype 2 Strain D39 of Streptococcus pneumoniae and Comparison with That of Unencapsulated Laboratory Strain R6. Journal of Bacteriology 189, 38–51 (2007).

46. Trouve, J. et al. Nanoscale dynamics of peptidoglycan assembly during the cell cycle of Streptococcus pneumoniae. Curr Biol 31, 2844–2856.e6 (2021).

47. Lamanna, M. M. et al. Roles of RodZ and class A PBP1b in the assembly and regulation of the peripheral peptidoglycan elongasome in ovoid-shaped cells of Streptococcus pneumoniae D39. Molecular Microbiology 118, 336–368 (2022).

48. Land, A. D. & Winkler, M. E. The Requirement for Pneumococcal MreC and MreD Is Relieved by Inactivation of the Gene Encoding PBP1a. Journal of Bacteriology 193, 4166–4179 (2011).

49. Claessen, D. et al. Control of the cell elongation–division cycle by shuttling of PBP1 protein in Bacillus subtilis. Molecular Microbiology 68, 1029–1046 (2008).

50. Rismondo, J., Wamp, S., Aldridge, C., Vollmer, W. & Halbedel, S. Stimulation of PgdA-dependent peptidoglycan N-deacetylation by GpsB-PBP A1 in Listeria monocytogenes. Molecular Microbiology 107, 472–487 (2018).

51. Fleurie, A. et al. Interplay of the serine/threonine-kinase StkP and the paralogs DivIVA and GpsB in pneumococcal cell elongation and division. PLoS Genet 10, e1004275 (2014).

52. Pompeo, F., Foulquier, E., Serrano, B., Grangeasse, C. & Galinier, A. Phosphorylation of the cell division protein GpsB regulates PrkC kinase activity through a negative feedback loop in acillus subtilis. Molecular Microbiology 97, 139–150 (2015).

53. Halbedel, S. & Lewis, R. J. Structural basis for interaction of DivIVA/GpsB proteins with their ligands. Molecular Microbiology 111, 1404–1415 (2019).

54. Trouve, J. et al. DivIVA controls the dynamics of septum splitting and cell elongation in Streptococcus pneumoniae. mBio 0, e01311–24 (2024).

55. Berg, K. H., Biørnstad, T. J., Straume, D. & Håvarstein, L. S. Peptide-Regulated Gene Depletion System Developed for Use in Streptococcus pneumoniae. Journal of Bacteriology 193, 5207–5215 (2011).

56. Sung, C. K., Li, H., Claverys, J. P. & Morrison, D. A. An rpsL cassette, janus, for gene replacement through negative selection in Streptococcus pneumoniae. Appl Environ Microbiol 67, 5190–5196 (2001).

57. Zucchini, L. et al. PASTA repeats of the protein kinase StkP interconnect cell constriction and separation of Streptococcus pneumoniae. Nat Microbiol 3, 197–209 (2018).

58. Ducret, A., Quardokus, E. M. & Brun, Y. V. MicrobeJ, a tool for high throughput bacterial cell detection and quantitative analysis. Nat Microbiol 1, 1–7 (2016).

59. Morlot, C., Zapun, A., Dideberg, O. & Vernet, T. Growth and division of Streptococcus pneumoniae: localization of the high molecular weight penicillin-binding proteins during the cell cycle. Molecular Microbiology 50, 845–855 (2003).

60. Fleurie, A. et al. Mutational dissection of the S/T-kinase StkP reveals crucial roles in cell division of Streptococcus pneumoniae. Molecular Microbiology 83, 746–758 (2012).

61. Morlot, C. et al. Structure of the essential peptidoglycan amidotransferase MurT/GatD complex from Streptococcus pneumoniae. Nat Commun 9, 3180 (2018).

62. Breukink, E. et al. Lipid II Is an Intrinsic Component of the Pore Induced by Nisin in Bacterial Membranes *. Journal of Biological Chemistry 278, 19898–19903 (2003).

63. Helassa, N., Vollmer, W., Breukink, E., Vernet, T. & Zapun, A. The membrane anchor of penicillin-binding protein PBP2a from Streptococcus pneumoniae influences peptidoglycan chain length. The FEBS Journal 279, 2071–2081 (2012).

64. Barrett, D. et al. Analysis of glycan polymers produced by peptidoglycan glycosyltransferases. J Biol Chem 282, 31964–31971 (2007).

