## Supplemental Information for "Characterization of a GpsB-associated regulator of PBP1a reveals the organization of the cell wall remodeling complex of *Streptococcus pneumoniae*"

This file contains the legend of Supplementary Video 1 and Supplementary Tables.

**Supplementary Video 1: AF3 model of PBP1a in complex with GarP.**

The movie shows the conformation change in the interdomain of PBP1a upon the binding of GarP. PBP1a is in grey and GarP in pink.

**Table S1.** Strains and plasmids used in this study.

| Strains | Genotype and description | References | #primers (Table S2) |
|---|---|---|---|
| ***S. pneumoniae* strains** | | | |
| R800 | *S. pneumoniae* R6 derivative strain | Gift from JP Claverys (Toulouse-France) | |
| WT | R800 *rpsL1* ; Str$^{R}$ | *Fleurie et al., 2012*[60] | |
| *ΔgarP::kan-rpsL* | R800 *rpsL1, ΔgarP::kan-rpsL* ; Str$^{S}$ Kan$^{R}$ | This study | 1, 2, 3, 4, 5 and 6 |
| *ΔgarP* | R800 *rpsL1, ΔgarP* ; Str$^{R}$ | This study | 1, 2, 7 and 8 |
| *flag-garP* | R800 *rpsL1, garP::flag-garP* ; Str$^{R}$ | This study | 1, 2, 9, 10, 11 and 12 |
| *pComX-flag-garP* | R800 *rpsL1, ΔIS1167::P1::PcomR-comR, cpsN-O::pcomX-flag-garP* ; Str$^{R}$ | This study | 11, 12, 13, 14, 15 and 16 |
| *pComX-flag-garP, ΔgarP::kan-rpsL* | R800 *rpsL1, ΔIS1167::P1::PcomR-comR, cpsN-O::pcomX-flag-garP, ΔgarP::kan-rpsL* ; Str$^{S}$ Kan$^{R}$ | This study | 1 and 2 |
| *pComX-Flag-garP, ΔgarP* | R800 *rpsL1, ΔIS1167::P1::PcomR-comR, cpsN-O::pcomX-flag-garP, ΔgarP* ; Str$^{R}$ | This study | 1 and 2 |
| *Membrane probe* | R800 *rpsL1, cbpg-truncation::pEno-gfp-TM-Ex1$_{MapZ}$-cm* ; str$^{R}$ Cm$^{R}$ | This study | 17, 18, 19, 20, 21, 22, 23, 24, 25 and 26 |
| *Membrane probe, ΔgarP::kan-rpsL* | R800 *rpsL1, cbpg-truncation::pEno-gfp-TM-Ex1$_{MapZ}$-cm, ΔgarP::kan-rpsL* ; Str$^{S}$ Kan$^{R}$ Cm$^{R}$ | This study | 1 and 2 |
| *Membrane probe, ΔgarP* | R800 *rpsL1, cbpg-truncation::pEno-gfp-TM-Ex1$_{MapZ}$-cm ΔgarP* ; Str$^{R}$ Cm$^{R}$ | This study | 1 and 2 |
| *ftsZ-gfp* | R800 *rpsL1*, *ftsZ::ftsZ-gfp* ; Str$^{R}$ | *Fleurie et al., 2014*[51] | |
| *ftsZ-rfp* | R800 *rpsL1*, *ftsZ::ftsZ-rfp* ; Str$^{R}$ | *Fleurie et al., 2014*[51] | |
| *ftsZ-rfp, ΔgarP::kan-rpsL* | R800 *rpsL1*, *ftsZ::ftsZ-rfp, ΔgarP::kan-rpsL* ; Str$^{S}$ Kan$^{R}$ | This study | 1 and 2 |
| *ftsZ-rfp, ΔgarP* | R800 *rpsL1*, *ftsZ::ftsZ-rfp, ΔgarP* ; Str$^{R}$ | This study | 1 and 2 |
| *gfp-pbp2b* | R800 *rpsL1*, *pbp2b::gfp-pbp2b* ; Str$^{R}$ | *Fleurie et al., 2014*[51] | |
| *gfp-pbp2b, ΔgarP::kan-rpsL* | R800 *rpsL1*, *pbp2b::gfp-pbp2b, ΔgarP::kan-rpsL* ; Str$^{S}$ Kan$^{R}$ | This study | 1 and 2 |
| *gfp-pbp2b, ΔgarP* | R800 *rpsL1*, *pbp2b::gfp-pbp2b, ΔgarP* ; Str$^{R}$ | This study | 1 and 2 |

| | | | |
|---|---|---|---|
| gfp-pbp2x | R800 rpsL1, pbp2x::gfp-pbp2x ; $Str^R$ | Fleurie et al., 2014[51] | |
| gfp-pbp2x, ΔgarP::kan-rpsL | R800 rpsL1, pbp2x::gfp-pbp2x, ΔgarP::kan-rpsL ; $Str^S$ $Kan^R$ | This study | 1 and 2 |
| gfp-pbp2x, ΔgarP | R800 rpsL1, pbp2x::gfp-pbp2x, ΔgarP ; $Str^R$ | This study | 1 and 2 |
| gfp-mapZ | R800 rpsL1, mapZ::gfp-mapZ ; $Str^R$ | Fleurie et al., 2014[28] | |
| ΔgarP, ΔmapZ::kan-rpsL | R800 rpsL1, ΔgarP, ΔmapZ::kan-rpsL ; $Str^S$ $Kan^R$ | This study | 27 and 28 |
| gfp-mapZ, ΔgarP | R800 rpsL1, mapZ::gfp- mapZ, ΔgarP ; $Str^R$ | This study | 27 and 28 |
| gfp-garP | R800 rpsL1, garP::gfp-garP ; $Str^R$ | This study | 1, 2, 12, 29, 30 and 31 |
| garP-ΔlysM | R800 rpsL1, garP::garPΔ110-158 ; $Str^R$ | This study | 1, 2, 32 and 33 |
| garP-Δcyto | R800 rpsL1, garP::garPΔ2-31 ; $Str^R$ | This study | 1, 2, 9 and 34 |
| garP-ΔPαH | R800 rpsL1, garP::garPΔ61-76 ; $Str^R$ | This study | 1, 2, 35 and 36 |
| gfp-garP-ΔlysM | R800 rpsL1, garP::gfp-garPΔ110-158 ; $Str^R$ | This study | 1, 2, 32 and 33 |
| gfp-ΔgarP-Δcyto | R800 rpsL1, garP::gfp-garPΔ2-31 ; $Str^R$ | This study | 1, 2, 31 and 37 |
| gfp-garP-ΔPαH | R800 rpsL1, garP::gfp-garPΔ61-76 ; $Str^R$ | This study | 1, 2, 35 and 36 |
| gfp-garP-ΔcytoΔlysM | R800 rpsL1, garP::gfp-garPΔ2-31/Δ110-158 ; $Str^R$ | This study | 1, 2, 32 and 33 |
| gfp-garP-Δout | R800 rpsL1, garP::gfp-garPΔ58-158 ; $Str^R$ | This study | 1, 2, 32 and 38 |
| gfp-garP-ΔcytoΔout | R800 rpsL1, garP::gfp-garPΔ2-31/Δ58-158 ; $Str^R$ | This study | 1, 2, 32 and 38 |
| Δpbp1a::kan-rpsL | R800 rpsL1, Δpbp1a::kan-rpsL ; $Str^S$ $Kan^R$ | This study | 3, 4, 39, 40, 41 and 42 |
| Δpbp1a | R800 rpsL1, Δpbp1a ; $Str^R$ | This study | 39, 40, 43 and 44 |
| Δpbp2a::kan-rpsL | R800 rpsL1, Δpbp2a::kan-rpsL ; $Str^S$ $Kan^R$ | This study | 3, 4, 45, 46, 47 and 48 |
| Δpbp2a | R800 rpsL1, Δpbp2a ; $Str^R$ | This study | 45, 46, 49 and 50 |
| Δpbp1b::kan-rpsL | R800 rpsL1, Δpbp1b::kan-rpsL ; $Str^S$ $Kan^R$ | This study | 3, 4, 51, 52, 53 and 54 |
| Δpbp1b | R800 rpsL1, Δpbp1b ; $Str^R$ | This study | 51, 52, 55 and 56 |
| ΔmpgA::kan-rpsL | R800 rpsL1, ΔmpgA::kan-rpsL ; $Str^S$ $Kan^R$ | This study | 3, 4, 57, 58, 59 and 60 |
| ΔmacP::kan-rpsL | R800 rpsL1, ΔmacP::kan-rpsL ; $Str^S$ $Kan^R$ | This study | 3, 4, 61, 62, 63 and 64 |
| ΔgpsB::kan-rpsL | R800 rpsL1, ΔgpsB::kan-rpsL ; $Str^S$ $Kan^R$ | This study | 3, 4, 65, 66, 67 and 68 |
| ΔcbpM::kan-rpsL | R800 rpsL1, ΔcbpM::kan-rpsL ; $Str^S$ $Kan^R$ | This study | 3, 4, 69, 70, 71 and 72 |
| gfp | R800 rpsL1, ΔIS1167:: pCEP-gfp ; $Str^R$ $Kan^R$ | This study | 73, 74, 75 and 76 |
| garP-PαH-$R_{64}$A | R800 rpsL1, garP::garP-R64A ; $Str^R$ | This study | 1, 2, 77 and 78 |
| garP-PαH-$T_{65}$A | R800 rpsL1, garP::garP-T65A ; $Str^R$ | This study | 1, 2, 79 and 80 |
| garP-PαH-$L_{68}$A | R800 rpsL1, garP::garP-L68A ; $Str^R$ | This study | 1, 2, 81 and 82 |
| garP-PαH-$K_{69}$A | R800 rpsL1, garP::garP-K69A ; $Str^R$ | This study | 1, 2, 83 and 84 |
| garP-PαH-$D_{70}$A | R800 rpsL1, garP::garP-D70A ; $Str^R$ | This study | 1, 2, 85 and 86 |
| garP-PαH-$F_{71}$A | R800 rpsL1, garP::garP-F71A ; $Str^R$ | This study | 1, 2, 87 and 88 |
| garP-PαH-$H_{72}$A | R800 rpsL1, garP::garP-H72A ; $Str^R$ | This study | 1, 2, 89 and 90 |
| garP-PαH-$D_{73}$A | R800 rpsL1, garP::garP-D73A ; $Str^R$ | This study | 1, 2, 91 and 92 |
| garP-PαH-$S_{74}$A | R800 rpsL1, garP::garP-S74A ; $Str^R$ | This study | 1, 2, 93 and 94 |
| garP-PαH-$D_{75}$A | R800 rpsL1, garP::garP-D75A ; $Str^R$ | This study | 1, 2, 95 and 96 |
| garP-PαH-$L_{68}$A/$F_{71}$A | R800 rpsL1, garP::garP-L68A/F71A ; $Str^R$ | This study | 1, 2, 97 and 98 |
| pbp1a-$A_{124}$T | R800 rpsL1, pbp1a:: pbp1a -A124T ; $Str^R$ | This study | 39 and 40 |

| | | | |
|---|---|---|---|
| Δ*garP*, Δ*pbp1a::kan-rpsL* | R800 *rpsL1*, Δ*garP*, Δ*pbp1a::kan-rpsL* ; $Str^S$ $Kan^R$ | This study | 39 and 40 |
| *pbp1a-A*$_{124}$*T*, Δ*garP* | R800 *rpsL1, pbp1a:: pbp1a -A124T*, Δ*garP* ; $Str^R$ | This study | 39 and 40 |
| *garP-P*α*H-L*$_{68}$*A/F*$_{71}$*A*, Δ*pbp1a::kan-rpsL* | R800 *rpsL1, garP::garP-L68A/F71A*, Δ*pbp1a::kan-rpsL* ; $Str^S$ $Kan^R$ | This study | 39 and 40 |
| *pbp1a-A*$_{124}$*T, garP-P*α*H-L*$_{68}$*A/F*$_{71}$*A* | R800 *rpsL1, pbp1a:: pbp1a -A124T, garP::garP-L68A/F71A* ; $Str^R$ | This study | 39 and 40 |
| Δ*pgdA::kan-rpsL* | R800 *rpsL1*, Δ*pgdA::kan-rpsL* ; $Str^S$ $Kan^R$ | This study | 3, 4, 99, 100, 101 and 102 |
| Δ*pgdA* | R800 *rpsL1*, Δ*pgdA* ; $Str^R$ | This study | 99, 100, 103 and 104 |
| ***E. coli* strains** | | | |
| *XL1-Blue* | *supE44 hsdR17 recA1 endA1 gyrA46 thi relA1 lac-F'[proAB + lacI$^q$lacZ*Δ*M15 Tn10 (Tc$^R$)]* | Bullock et al. 1987 | |
| *TOP10* | *F- mcrA Δ(mrr-hsdRMS-mcrBC) φ80lacZΔM15 ΔlacX74 recA1 araD139 Δ(ara-leu)7697 galU galK rpsL (StrR) endA1 nupG* | Invitrogen | |
| *DH5*α | *F– φ80lacZΔ M15 Δ (lacZYA-argF) U169 recA1 endA1 hsdR17 (rK– mK+) PhoA supE44 λ- thi–1 gyrA96 relA1* | Lucigen | |
| *BTH101* | *F-, cya-99, araD139, galE15, galK16, rpsL1 (Str$^R$), hsdR2, mcrA1, mcrB1* | Euromedex | |
| *BL21 (DE3)* | *F-, ompT gal dcm lon hsdS$_B$ (r$_B$-m$_B$-) λ(DE3 [lacI lacUV5-T7 gene 1 ind1 sam7 nin5])* | Studier et al. 1986 | |
| ***Plasmids*** | | | |
| *pt7.7-garP*$_{full\ length}$*-TEV-6His* | *pt7.7-TEV-6His derivative, encoding GarP from Met1 to Lys158* ; $Amp^R$ | This study | 105, 106, 107 and 108 |
| *pET46-pbp1a* | *pET46(LIC) derivative, encoding PBP1a from Val31 to Pro719 ;* $Amp^R$ | Zapun et al. 2013[34] | |
| *pKNT25* | *pSU40 derivative, encoding T25 fragment of the CyaA B. pertussis protein for C-terminal T25 fusion ;* $Kan^R$ | Karimova et al., 1998 | |
| *pUT18* | *pUC19 derivative, encoding T18 fragment of the CyaA B. pertussis protein for C-terminal T18 fusions ;* $Amp^R$ | Karimova et al., 1998 | |
| *pKT25-zip* | *pKT25 derivative, encoding T25 fragment, fused to the leucine zipper region of the GCN4 yeast protein ;* $Kan^R$ | Karimova et al., 1998 | |
| *pUT18C-zip* | *pUT18C derivative, encoding T18 fragment, fused to the leucine zipper region of the GCN4 yeast protein ;* $Amp^R$ | Karimova et al., 1998 | |
| *pKT25-pbp1a* | *pKT25 derivative, encoding PBP1a from Met1 to , fused to T25 ;* $Kan^R$ | This study | 109, 110, 111 and 112 |
| *pUT18C-garP*$_{full\ length}$ | *pUT18C derivative, encoding GarP from Met1 to Lys158 fused to T18 ;* $Kan^R$ | This study | 113, 114, 115 and 116 |
| *pUT18C-garP-*Δ*cyto* | *pUT18C-garP*$_{full\ length}$ *derivative, encoding GarP from Met31 to Lys158 fused to T18 ;* $Kan^R$ | This study | 114 and 117 |
| *pUT18C-garP-*Δ*lysM* | *pUT18C-garP*$_{full\ length}$ *derivative, encoding GarP from Met1 to Gly110 fused to T18 ;* $Kan^R$ | This study | 113 and 118 |
| *pUT18C-garP-*Δ*lysM*Δ*cyto* | *pUT18C-garP-*Δ*lysM derivative, encoding GarP from Met31 to Gly110 fused to T18 ;* $Kan^R$ | This study | 114 and 117 |

| | | | |
|---|---|---|---|
| *pUT18C-garP-ΔPαH* | *pUT18C-garP$_{full\ length}$ derivative, encoding GarP from Met1 to Lys158 with deletion of Gly61-Ala76 fused to T18 ;* $Kan^R$ | This study | 35 and 36 |
| *pUT18C-garP-ΔPαHΔcyto* | *pUT18C-garP-Δcyto derivative, encoding GarP from Met31 to Lys158 with deletion of Gly61-Ala76 fused to T18 ;* $Kan^R$ | This study | 35 and 36 |
| *pUT18C-garP-ΔlysMΔPαHΔcyto* | *pUT18C-garP-ΔlysMΔcyto derivative, encoding GarP from Met31 to Gly110 with deletion of Gly61-Ala76 fused to T18 ;* $Kan^R$ | This study | 35 and 36 |
| *pUT18-gpsB* | *pUT18 derivative, encoding GpsB from Met1 to Phe113, fused to T18 ;* $Amp^R$ | *Fleurie et al., 2014*[51] | |
| *pKNT25-gpsB* | *pKNT25 derivative, encoding GpsB from Met1 to Phe113, fused to T25 ;* $Kan^R$ | *Fleurie et al., 2014*[51] | |
| *pUT18C-gpsB* | *pUT18C derivative, encoding GpsB from Met1 to Phe113, fused to T18 ;* $Amp^R$ | *Fleurie et al., 2014*[51] | |
| *pKT25-gpsB* | *pKT25 derivative, encoding GpsB from Met1 to Phe113, fused to T25 ;* $Kan^R$ | *Fleurie et al., 2014*[51] | |
| *pUT18C-garPΔmotif* | *pUT18C-garP$_{full\ length}$ derivative, encoding GarP from Met1 to Lys158 with deletion of Ser16-Arg21 fused to T18 ;* $Kan^R$ | This study | 119 and 120 |
| *pUT18C-pgdA* | *pUT18C derivative, encoding PgdA from Met1 to Glu463, fused to T18 ;* $Amp^R$ | This study | 113, 114, 121 and 122 |
| *pUT18C-pgdAΔmotif* | *pUT18C derivative, encoding PgdA from Met1 to Glu463 with deletion of Ser4-Arg8, fused to T18 ;* $Amp^R$ | This study | 114 and 123 |
| *pUT18C-pbp2a* | *pUT18C derivative, encoding PBP2a from Met1 to Arg731, fused to T18 ;* $Amp^R$ | This study | 113, 114, 124 and 125 |
| *pUT18C-pbp2aΔmotif* | *pUT18C derivative, encoding PBP2a from Met1 to Arg731 with deletion of Ser32-Lys37, fused to T18 ;* $Amp^R$ | This study | 126 and 127 |
| *pKT25-mpgA* | *pKT25 derivative, encoding MpgA from Leu1 to Asn551, fused to T25 ;* $Kan^R$ | This study | 109, 110, 128 and 129 |
| *pKT25-mpgAΔmotif* | *pKT25 derivative, encoding MpgA from Leu1 to Asn551 with deletion of Ser165-Arg170, fused to T25 ;* $Kan^R$ | This study | 130 and 131 |

**Table S2.** Primers used in this study.

| # | Primer name | +/- | Sequence 5’-3’ |
|---|---|---|---|
| 1 | *Upstream region of garP* | + | AGCTCTGCCGTCGGGTGTCC |
| 2 | *Downstream region of garP* | - | CCTTATCATCATTCACTATT |
| 3 | *5'- [kan-rpsL]* | + | CCGTTTGATTTTTAATGGATAATG |
| 4 | *3'- [kan-rpsL]* | - | AGAGACCTGGGCCCCTTTCC |
| 5 | *[kan-rpsL]-Downstream region of garP* | + | CATAAGGAAAGGGGCCCAGGTCTCTTCGTGCAGGAATCTCCATTG |
| 6 | *[kan-rpsL]-Upstream region of garP* | - | CATTATCCATTAAAAATCAAACGGACCTTACTCCTTGTTT |
| 7 | *5’-deletion-garP* | + | ATGGCAAAAGAACCGTGGCAAGGCCTTGAATCCTGGGCACAT |
| 8 | *3’-deletion-garP* | - | ATGTGCCCAGGATTCAAGGCCTTGCCACGGTTCTTTTGCCAT |
| 9 | *Upstream of garP* | - | ACCTTACTCCTTGTTTTTTTTTAC |
| 10 | *Upstream of garP- FLAG* | + | AGTAAAAAAAAACAAGGAGTAAGGTATGGACTACAAAGACCATGACG |
| 11 | *FLAG-linker2* | - | CTGCAGGAACTCGATGTCTAGTTTTTTATCGTCGTCATCTTTGTAGTC |
| 12 | *Linker2-garP* | + | AAACTAGACATCGAGTTCCTGCAGATGGCAAAAGAACCGTGGCA |
| 13 | *Upstream region of pComX platform* | + | ATAACAAATCCAGTAGCTTTGG |
| 14 | *garP-downstream region of pComX platform* | - | ATTGGGAAGAGTTACATATTAGAAACTATTTTATTTTTATAACAT |
| 15 | *Downstream region of pComX platform* | - | CATCGGAACCTATACTCTTTTAG |
| 16 | *Downstream of pComX promoter* | + | TTTCTAATATGTAACTCTTCCCAAT |
| 17 | *Upstream region of cbpG*$_{truncation}$ | + | TACTCTTCGAAAATCTCTTC |
| 18 | *Downstream region of cbpG*$_{truncation}$ | - | TCTGTTGATGCGAGAATCAA |
| 19 | *pEno* | + | GAAAACAGTATATCATAAAA |
| 20 | *cbpG*$_{truncation}$*-pEno* | - | TTTTATGATATACTGTTTTCGTAAAGTATTATCAATTTTA |
| 21 | *pEno* | - | TTTTTACTCTCCTTATGAGTTAAATTTTTTACACC |
| 22 | *pEno-GFP* | + | CTCATAAGGAGAGTAAAAAATGATTTCTAAAGGTGAAGAATTG |
| 23 | *Ex1*$_{mapZ}$*-Cm*$^{R}$ | - | ACTAGTGCTCATTAGAATTCTTAACTTCTAGTCTCATTTG |
| 24 | *Upstream-Cm*$^{R}$ | + | AAGAATTCTAATGAGCACT |
| 25 | *Downstream-Cm*$^{R}$ | - | TTATAAAAGCCAGTCATTAG |
| 26 | *CmR-cbpG*$_{truncation}$ | + | CTAATGACTGGCTTTTATAAATCAAATTAACAGTGCAGTTA |

| | | | |
|---|---|---|---|
| 27 | *Upstream region of garP* | + | GTCTAGCCTCTTTAAACGTGG |
| 28 | *Downstream region of garP* | - | GCATCAAGTCATAGCTTTCTGC |
| 29 | *GFPstart* | + | ATGATTTCTAAAGGTGAAGAATT |
| 30 | *Upstream of garP-GFP* | - | ATTCTTCACCTTTAGAAATCATACCTTACTCCTTGTTTTTTTTTA |
| 31 | *GFP-linker2* | - | CTGCAGGAACTCGATGTCTAGTTTTTATACAATTCATCCATACCATGTG |
| 32 | *Downstream of garP* | + | GAGCCGATATGAAAACAAT |
| 33 | *ΔlysM of garP* | - | GAATTGTTTTCATATCGGCTCCTATCCTTCAGGATTAGCAGC |
| 34 | *Δcyto of garP* | + | GTAAAAAAAAACAAGGAGTAAGGTATGGCTAATCGTATTTTGACG |
| 35 | *5’-ΔPαH of GarP* | + | TCTCATCTATCTATCATCGGGGAGTGTAGTACAAATCTCATC |
| 36 | *3’-ΔPαH of GarP* | - | CCCCGATGATAGATAGATGAG |
| 37 | *Gfp-garP-Δcyto* | + | AAACTAGACATCGAGTTCCTGCAGATGGCTAATCGTATTTTGAC |
| 38 | *garP-Δout* | - | GAATTGTTTTCATATCGGCTCCTATAGATAGATGAGAACG |
| 39 | *Upstream region of pbp1a* | + | AAGCAGAATAGCCTA |
| 40 | *Downstream region of pbp1a* | - | GATTTTGGTAAGACAATTAAGGC |
| 41 | *[kan-rpsL]-Downstream region of pbp1a* | + | GGAAAGGGGCCCAGGTCTCTCATTTATCATCCAGATTTTTCTG |
| 42 | *[kan-rpsL]-Upstream region of pbp1a* | - | CATTATCCATTAAAAATCAAACGGTCATCTTGTTTTACCACCTAATAAATG |
| 43 | *Upstream of pbp1a* | - | TCATCTTGTTTTACCACCTAATAAATG |
| 44 | *5’-deletion-pbp1a* | + | CATTTATTAGGTGGTAAAACAAGATGACATTTATCATCCAGATTTTTCTG |
| 45 | *Upstream region of pbp2a* | + | CTACGATCATGGCGATCACG |
| 46 | *Downstream region of pbp2a* | - | CAAGAACATAACCTGGAAAGCG |
| 47 | *[kan-rpsL]-Downstream region of pbp2a* | + | GGAAAGGGGCCCAGGTCTCTGATGCTTGTCAAAGCCTAGC |
| 48 | *[kan-rpsL]-Upstream region of pbp2a* | - | CATTATCCATTAAAAATCAAACGGGCGTTTATTTTATCATCTTCATC |
| 49 | *Upstream of pbp2a* | - | GCGTTTATTTTATCATCTTCATC |
| 50 | *5’-deletion-pbp2a* | + | GATGAAGATGATAAAATAAACGCGATGCTTGTCAAAGCCTA |
| 51 | *Upstream region of pbp1b* | + | CCCTTGCATGATTTGGTAAGC |

| | | | |
|---|---|---|---|
| 52 | *Downstream region of pbp1b* | - | GCGAGAGGACATTGCTAAAC |
| 53 | *[kan-rpsL]-Downstream region of pbp1b* | + | GGAAAGGGGCCCAGGTCTCTTTCTCTAAATGAAGTGGCCAATC |
| 54 | *[kan-rpsL]-Upstream region of pbp1b* | - | CATTATCCATTAAAAATCAAACGGGGATTTCCTCACTTTATCTATTATACC |
| 55 | *Upstream of pbp1b* | - | GGATTTCCTCACTTTATCTATTATACC |
| 56 | *5'-deletion-pbp1b* | + | GGTATAATAGATAAAGTGAGGAAATCCACAGGTTCGACTGTTACCAG |
| 57 | *Upstream region of mpgA* | + | CCTTGAAAGAGGTGGCAGTTGAG |
| 58 | *Downstream region of mpgA* | - | TTAGGCTGTTTTTTCAACCTTCAAGAT |
| 59 | *[kan-rpsL]-Downstream region of mpgA* | + | GGAAAGGGGCCCAGGTCTCACAAACTAAAATTATGTGATAC |
| 60 | *[kan-rpsL]-Upstream region of mpgA* | - | CATTATCCATTAAAAATCAAACGGTTCACTCAAAAGTTTTTCCTCC |
| 61 | *Upstream region of macP* | + | ATGGAGTTTGAAGAAAAAACGCTTAGCC |
| 62 | *Downstream region of macP* | - | TTAATCTAAAGCCTTCAAAAAGGCC |
| 63 | *[kan-rpsL]-Downstream region of macP* | + | GGAAAGGGGCCCAGGTCTCTTAGAAAAGGAATTGAAATG |
| 64 | *[kan-rpsL]-Upstream region of macP* | - | CCATTAAAAATCAAACGGACTGACCTCCTCTATTTTTTCTG |
| 65 | *Upstream region of gpsB* | + | GCCAAGCCCTGAGACAAATAG |
| 66 | *Downstream region of gpsB* | - | GGAACCGAGCTCCAAGTGG |
| 67 | *[kan-rpsL]-Downstream region of gpsB* | + | GGAAAGGGGCCCAGGTCTCTGTAGTTATTTGAGATGTGCAATTTTTGG |
| 68 | *[kan-rpsL]-Upstream region of gpsB* | - | CATTATCCATTAAAAATCAAACGGTCTCGCTTGCTAGTATTATTATATAAAAAAAGGC |
| 69 | *Upstream region of cbpM* | + | TCAGAGATGATACTAGAG |
| 70 | *Downstream region of cbpM* | - | AGTGGAAGGTCGTGGATGCC |

| | | | |
|---|---|---|---|
| 71 | *[kan-rpsL]-Downstream region of cbpM* | + | GGAAAGGGGCCCAGGTCTCTTAGATGGCTATAGAGTCAA |
| 72 | *[kan-rpsL]-Upstream region of cbpM* | - | ATCCATTAAAAATCAAACGGCTCTTTAGTAATTTGACGAG |
| 73 | *Upstream region of IS1167$_{truncation}$* | + | ACCCTCATGAATTCTCAGGCGGT |
| 74 | *Downstream region of IS1167$_{truncation}$* | - | ATGAAGAAATACCAACAATTATTTAAGCAAATCC |
| 75 | *5'-GFP-pUC57* | + | ATGAATTCCCATGGTTTCTAAAGGTG |
| 76 | *3'GFP-pUC57* | - | TATGGATCCTTATTTATACAATTCATCCATACCATGTG |
| 77 | *5'-garP-R64A* | + | CATCGGGGGGGAGTAATGCCACAGCAGCCTTAAAAGAC |
| 78 | *3'-garP-R64A* | - | GTCTTTTAAGGCTGCTGTGGCATTACTCCCCCCCGATG |
| 79 | *5'-garP-T65A* | + | CGGGGGGGAGTAATCGCGCAGCAGCCTTAAAAGACTTTC |
| 80 | *3'-garP-T65A* | - | GAAAGTCTTTTAAGGCTGCTGCGCGATTACTCCCCCCCG |
| 81 | *5'-garP-L68A* | + | GTAATCGCACAGCAGCCGCAAAAGACTTTCATGATTCTGATG |
| 82 | *3'-garP-L68A* | - | CATCAGAATCATGAAAGTCTTTTGCGGCTGCTGTGCGATTAC |
| 83 | *5'-garP-K69A* | + | GTAATCGCACAGCAGCCTTAGCAGACTTTCATGATTCTGATGC |
| 84 | *3'-garP-K69A* | - | GCATCAGAATCATGAAAGTCTGCTAAGGCTGCTGTGCGATTAC |
| 85 | *5'-garP-D70A* | + | GCACAGCAGCCTTAAAAGCCTTTCATGATTCTGATGCAAG |
| 86 | *3'-garP-D70A* | - | CTTGCATCAGAATCATGAAAGGCTTTTAAGGCTGCTGTGC |
| 87 | *5'-garP-F71A* | + | CACAGCAGCCTTAAAAGACGCTCATGATTCTGATGCAAGTGTAG |
| 88 | *3'-garP-F71A* | - | CTACACTTGCATCAGAATCATGAGCGTCTTTTAAGGCTGCTGTG |
| 89 | *5'-garP-H72A* | + | CAGCCTTAAAAGACTTTGCTGATTCTGATGCAAGTGTAG |
| 90 | *3'-garP-H72A* | - | GTACTACACTTGCATCAGAATCAGCAAAGTCTTTTAAGGCTGC |
| 91 | *5'-garP-D73A* | + | GCAGCCTTAAAAGACTTTCATGCTTCTGATGCAAGTGTAGTACAAATC |
| 92 | *3'-garP-D73A* | - | GATTTGTACTACACTTGCATCAGAAGCATGAAAGTCTTTTAAGGCTGC |
| 93 | *5'-garP-S74A* | + | GCAGCCTTAAAAGACTTTCATGATGCTGATGCAAGTGTAGTACAAATCTC |
| 95 | *3'-garP-S74A* | - | GAGATTTGTACTACACTTGCATCAGCATCATGAAAGTCTTTTAAGGCTGC |
| 95 | *5'-garP-D75A* | + | CCTTAAAAGACTTTCATGATTCTGCTGCAAGTGTAGTACAAATCTCATCTT C |
| 96 | *3'-garP-D75A* | - | GAAGATGAGATTTGTACTACACTTGCAGCAGAATCATGAAAGTCTTTTAAG G |
| 97 | *5'-garP-L68A/F71A* | + | GTAATCGCACAGCAGCCGCAAAAGACGCTCATGATTCTGATGCAAGTGTA G |
| 98 | *3'-garP-L68A/F71A* | - | CTACACTTGCATCAGAATCATGAGCGTCTTTTGCGGCTGCTGTGCGATTA C |
| 99 | *Upstream region of pgdA* | + | TCTATACTAGTATAGACGTCTTGGC |
| 100 | *Downstream region of pgdA* | - | CGCATTGCGAGTTTTCCATGC |
| 101 | *[kan-rpsL]-Downstream region of pgdA* | + | AGGAAAGGGGCCCAGGTCTCTGCAAGAAAAAATAGGTCTGTTAG |
| 102 | *[kan-rpsL]-Upstream region of pgdA* | - | CATTATCCATTAAAAATCAAACGGATTTATATCATAACAAATTTGAGCAATAA TTTCAATG |
| 103 | *Upstream of pgdA* | + | GCAAGAAAAAATAGGTCTGTTAGATATTTG |
| 104 | *3'-deletion-pgdA* | - | CAAATATCTAACAGACCTATTTTTTCTTGCATTTATATCATAACAAATTTGAG CAATAATTTC |
| 105 | *5'-Pt7.7* | + | CTGCAGGAAAACTTGTATTTCCAGG |
| 106 | *3'-Pt7.7* | - | CATATGTATATCTCCTTCTTAAAGTTAAAC |

| | | | |
|---|---|---|---|
| 107 | *5’-Pt7.7-garP* | + | CTTTAAGAAGGAGATATACATATGGCAAAAGAACCGTGGCAAG |
| 108 | *3’-Pt7.7-garP* | - | AATACAAGTTTTCCTGCAGTTTTATTTTTATAACATCACC |
| 109 | *5’-pKT25* | + | CGGGTACCTAAGTAACTAAGAATTC |
| 110 | *3’-pKT25* | - | CTCTAGAGTCGACCCTGCAG |
| 111 | *5’-pKT25-pbp1a* | + | TATTCTAGAGATGAACAAACCAACGATTCTG |
| 112 | *3’-pKT25-pbp1a* | - | TATGGTACCCGTGGTTGTGCTGGTTGAGGATTC |
| 113 | *5’-pUT18C* | + | CGGGTACCGAGCTCGAATTCA |
| 114 | *3’-pUT18C* | - | CTCTAGAGTCGACCTGCAG |
| 115 | *5’-pUT18C-garP* | + | ACTGCAGGTCGACTCTAGAGATGGCAAAAGAACCGTGG |
| 116 | *3’-pUT18C-garP* | - | TCGATGAATTCGAGCTCGGTACCCGTTTTATTTTTATAACATCAC |
| 117 | *5’-pUT18C-garP-∆cyto* | + | CCACTGCAGGTCGACTCTAGAGATGGCTAATCGTATTTTGAC |
| 118 | *3’-pUT18C-garP-∆lysM* | - | GAATTCGAGCTCGGTACCCGTCCTTCAGGATTAGCAGCTTC |
| 119 | *5’-garP-∆motif* | + | CATCGAAACCACGGAG |
| 120 | *3’-garP-∆motif* | - | CCCCTCCGTGGTTTCGATGTTCTTCTTGATCATAAATATC |
| 121 | *5’-pUT18C-pgdA* | + | CTAGTCTAGAGATGAATAAAAGTAGACTAGGACGTGG |
| 122 | *3’-pUT18C-pgdA* | - | TATGGTACCCGTTATTCATCACGACTATAGTACAGCTC |
| 123 | *5’-pUT18C-pgdA-∆motif* | + | CACTGCAGGTCGACTCTAGAGATGAATAAAGGCAGACACGGGAAAAC |
| 124 | *5’-pUT18C-pbp2a* | + | TATGGATCCCATGAAATTAGATAAATTATTTGAG |
| 125 | *3’-pUT18C-pbp2a* | - | TATGAGCTCTTAGCGAAATAGATTGACTATC |
| 126 | *5’-pbp2a-∆motif* | + | GATTCTACTATCTTACGTCGCAAATTAGCCCAAGTAGGTCC |
| 127 | *3’-pbp2a-∆motif* | - | GCGACGTAAGATAGTAGAATC |
| 128 | *5’-pKT25-mpgA* | + | CGGGCTGCAGGGTCGACTCTAGAGTTGAGTGAAAAGTCAAG |
| 129 | *3’-pKT25-mpgA* | - | CGAATTCTTAGTTACTTAGGTACCCGGTTTAATTTGCTGTTGAC |
| 130 | *5’-mpgA-∆motif* | + | CAGTGGATATCATCAGAGATACAGAAGGAGCAAAACCC |
| 131 | *3’-mpgA-∆motif* | - | TGTATCTCTGATGATATCCACTG |
